# Heterotypic interactions and sequence features modulate cellular reflectin condensate dynamics

**DOI:** 10.64898/2026.08.27.747642

**Authors:** Chi Phan, Rika Watanabe, Vinh Quang Le, Susan Walsh, Robert Levenson

**Affiliations:** Life Sciences Concentration, Soka University of America, Aliso Viejo, CA 92656, USA

**Keywords:** biophotonics, intrinsically disordered proteins, tunable, iridescence, protein self-assembly, biomolecular condensates, protein phase separation, reflectins

## Abstract

Reflectin proteins drive dynamic structural coloration in cephalopods by organizing into dense intracellular lamellar structures that dictate local refractive index. While reconstituted reflectins readily undergo liquid-liquid phase separation *in vitro*, these assemblies frequently undergo dynamic arrest, vitrifying into non-dynamic condensates. Here, we investigate how primary sequence features, post-translational modifications, and heterotypic interactions regulate the material properties of reflectin condensates within the environment of mammalian HeLa cells. Using confocal microscopy and fluorescence recovery after photobleaching (FRAP), we demonstrate that canonical block copolymeric A-type reflectins readily form dynamically arrested condensates, with the linker blocks primarily responsible for the observed arrest. In contrast, non-canonical B/C reflectin variants exhibit significantly greater fluidity and rapid recovery kinetics. We show that phosphomimetic substitutions progressively fluidize some reflectin condensates. Lastly, we find that heterotypic condensates composed of canonical and non-canonical reflectins in combinations associated with reversible iridescence in squid substantially enhance canonical mobility. Our findings establish a biophysical framework in which phosphorylation and heterotypic mixing cooperatively suppress dynamic arrest, enabling the reversible material transitions required for active cephalopod camouflage and communication.

## Introduction

Reflectins are specialized structural proteins that drive tunable biophotonics and dynamic structural coloration within specialized cephalopod cells called iridocytes or iridophores^1,2^. To fulfill their biological functions, these specialized cells maintain exceptionally high intracellular concentrations of reflectin proteins^3–6^. Reflectins are intrinsically disordered proteins (IDPs) characterized by a highly repetitive, block copolymer-like sequence architecture. This architecture consists of alternating conserved, highly ampholytic domains (Reflectin Motifs, RMs) and less-conserved, cationic linker regions^7–9^ (Fig. 1A). Reflectins have been investigated for their potential biotechnological applications, including as bioinspiration for novel materials^1,6,10,11^. Sequence analyses demonstrate that reflectins feature an unusual composition; they are heavily enriched in aromatic tyrosine and nonpolar methionine residues, and exhibit a near-complete absence of aliphatic hydrophobic residues (Fig. 1B). The unique amino acid composition and sequence architecture of reflectins position them well for analysis within a classic sticker-and-spacers framework, where multi-valent aromatic and electrostatic interactions (stickers) and non-interacting regions (spacers) govern condensate material properties^12,13^. In this model, both the number of interacting sites (chain valency) and the types of interactions between and within chains determine condensate microscopic and macroscopic properties.

**Figure 1.**
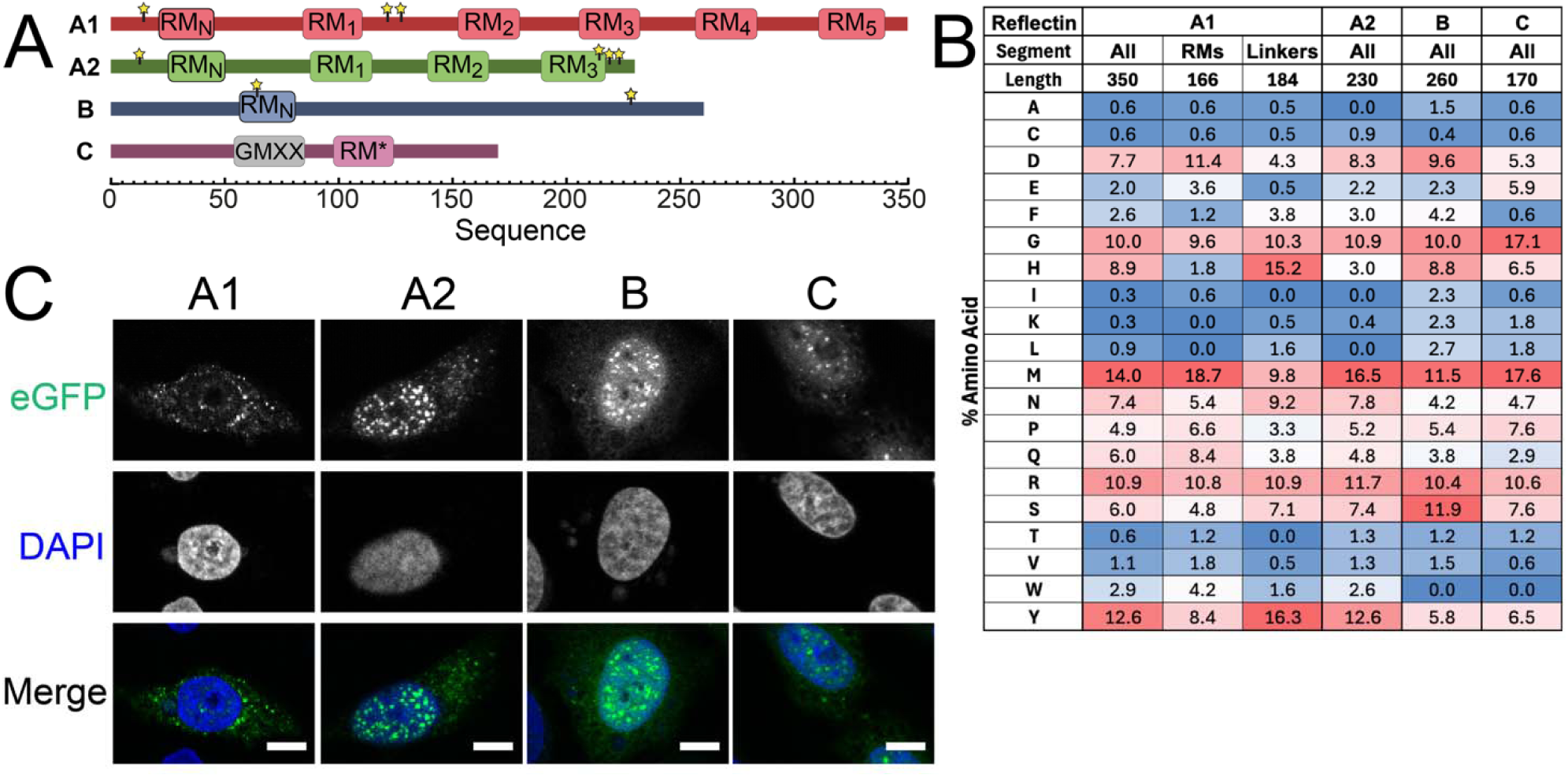
Schematics and expression of *D. opalescens* reflectins. A) Scaled linear schematics of *D. opalescens* reflectin isoforms A1, A2, B, and C. Narrow segments represent linkers, while wider segments indicate the conserved Reflectin Motifs (RMs). B) Compositional profile comparing selected residues and across the *D. opalescens* reflectins as well as the segments of A1. Compositional profile of all amino acids and linker and domain segments of all four *D. opalescens* proteins is provided in Fig. S1. C) Fixed cell confocal images of eGFP-labeled reflectin isoforms. Scale bar = *10* μm.

In the long-finned pelagic squid *Doryteuthis opalescens*, multiple distinct reflectin sub-types have been identified, categorized broadly into canonical (A1 and A2) and non-canonical (B and C) isoforms^3,4^. *In vivo*, *D. opalescens*, like other loliginid squid, exhibits acetylcholine-dependent tunability of its iridocytes^14,15^, where exposure to acetylcholine (ACh) triggers a signal transduction cascade culminating in phosphorylation of canonical A-type reflectins and dephosphorylation of non-canonical variants^3,4^. Furthermore, tissue-level profiling demonstrates that dynamic optical tunability across different tissue types is strongly correlated with the relative abundance of non-canonical B/C reflectin isoforms relative to canonical ones^4^. Despite significant characterization, the exact molecular mechanisms by which these multi-isoform systems cooperatively interact to drive tunable structural color remain poorly understood.

Recent *in vitro* investigations have demonstrated that purified reflectin proteins undergo liquid-liquid phase separation (LLPS), forming condensed droplets responsive to changes in pH and ionic strength^16–19^. However, a persistent feature of these homotypic reconstituted condensates is their acute propensity for rapid dynamic arrest, frequently vitrifying into non-dynamic states under low-ionic-strength or high pH conditions^16,17,20^. While *in vitro* assays with purified proteins are critical for isolating core thermodynamic phase boundaries of these proteins, these assays lack the cellular constraints and macromolecular crowding present in living cells. Though recent studies have confirmed that overexpressed reflectins can form spherical assemblies in human cell lines^21,22^, the internal viscoelastic properties and real-time macromolecular dynamics of these assemblies inside cell lines have not been systematically characterized.

To uncover the sequence-level design rules driving tunable reflectin condensation and resolve the role of multi-component networks in driving tunable iridescence, we transiently overexpressed wild-type and mutated variants of both canonical and non-canonical reflectins within a heterologous mammalian cell model, utilizing both single-component (homotypic) and double-component (heterotypic) expression conditions. Using fluorescence recovery after photobleaching (FRAP), our results provide biophysical insights into the material state of intracellular reflectin condensates, demonstrating how sequence valency, post-translational phosphorylation, and heterotypic mixing may interact to maintain fluidic biophotonic biomolecular networks in cephalopods.

## Experimental Procedures

### Plasmid construction and cloning

Linear gene fragments encoding different reflectin constructs were synthesized by either Integrated DNA Technologies (IDT) or Twist Biosciences, Inc. and then cloned into a linearized eGFP-C1 vector using manufacturer protocols from the NEBBuilder HiFi DNA Assembly kit (New England Biolabs). To create mScarlet-labeled reflectins, mScarlet was subcloned from pmScarlet_C1 to replace eGFP in the eGFP-C1 vector, and reflectin inserts were cloned into this vector using the above method. pmScarlet_C1 was a gift from Dorus Gadella (Addgene plasmid # 85042; http://n2t.net/addgene:85042; RRID:Addgene_85042)^23^. Successful assembly of all constructs was confirmed using whole plasmid sequencing (Eurofins, Inc.).

### Cell culture and transfection

For transfection, HeLa cells cultured in a 60 mm dish at >80% confluency were detached using trypsin-EDTA and resuspended in 1 mL of Dulbecco’s Modified Eagle’s medium (DMEM) (ThermoFisher Scientific) containing 10% fetal bovine serum (FBS) (ThermoFisher Scientific).

For fixed cell imaging, 200 µL of concentrated cell suspension was diluted into 2mL fresh DMEM in a 2 mL well containing a sterile coverslip. The cells were grown overnight in a 37°C incubator with 5% CO_2_ before transfection using Lipofectamine 3000 (Invitrogen). For each transfection, 2 µg plasmid DNA was mixed with 3 µL P3000 in 75 µL Opti-MEM, followed by the addition of 75 µL Opti-MEM containing 3 µL Lipofectamine 3000. The combined mixture was incubated for 15 minutes at room temperature before being added dropwise to the plated cells. Transfected cells were incubated for 8-12 hours, and then cells were washed with 2 mL PBS (ThermoFisher Scientific), fixed with 1 mL 100% cold methanol (4°C), and stored at –20 °C for at least 20 minutes. After removal of methanol, cells were washed three times with 2 mL PBS for 5 minutes and incubated in 500 μL DAPI/PBS for 15 minutes at room temperature. After removing DAPI/PBS, coverslips containing cells were washed twice with 2 mL PBS and placed on glass slides with Fluoromount-G (ThermoFisher Scientific).

For live cell imaging, 200 µL >80% confluent HeLa cells were diluted into 2 mL fresh DMEM in a 35 mm well with a 14 mm glass bottom micro-well (MatTek, Inc.). The cells were incubated overnight at 37°C in a humidified incubator with 5% CO prior to transfection and then transfected using the same protocol described above.

### Cell imaging and FRAP

Transfected HeLa cells were imaged using a confocal fluorescence microscope (Nikon ECLIPSE Ti2) at 600X using an oil-immersion objective and an environmental control chamber (37°C, 5% CO_2_). A 488-nm laser was used to excite the GFP-tagged reflectin, while a 561-nm laser was used for mScarlet-tagged reflectin. Three pre-bleach measurements were performed for each condensate. Bleaching was performed for 1 second with a laser power of 8, and recovery was generally monitored for 2 minutes and 30 seconds, with an 8-second interval between images. All quantitative parameters were derived from at least three biological replicates, with a minimum of 8 separate cells per replicate, except where noted for constructs with high levels of dynamic arrest.

### FRAP kinetic fitting and statistical analysis

Raw FRAP recovery curves were background-subtracted and full-scale normalized to pre-bleach intensity. Normalized recovery curves were fitted to a single-term exponential function:

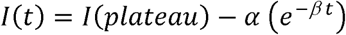

Fitting parameters were extracted to calculate mobile fraction (M_f_), and recovery half-time (t_1/2_) using equations:

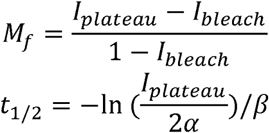

For full-scale normalized curves, *I_bleach_* = 0, so M_f_ = *I_plateau_*. M_f_ represents the fitted mobile fraction, and t_1/2_ is the recovery half-time. Single-term exponential function use was validated with an R^2^ ≥ 0.95 to average FRAP curves from single biological replicates, and Akaike Information Criterion (AIC) tests indicated that double-exponential functions resulted in parameter overfitting. Initial analysis of FRAP curves was performed using the easy-FRAP webserver^24^; final processing and analysis was performed using custom R scripts. For kinetic parameter extraction, individual recovery traces displaying R^2^ < 0.70 to a single-exponential model were removed from the dataset (4/1317 total measurements, excluding measurements removed for condensate drift or other technical issues). Traces exhibiting a fractional recovery of <15% by the end of the measurement period were classified as dynamically arrested and excluded from t_1/2_ calculations. For this population, M_f_ was quantified by raw fractional recovery at the final acquisition time point. To evaluate differences in M_f_ or *t*_1/2_ between constructs, a linear mixed model (LMM) was fitted to the cell-level data, where experimental replicate is treated as a random intercept to account for the hierarchical nesting of multiple cells within each independent biological replicate. Estimated marginal means (EMMs) were calculated to obtain adjusted group averages. Pairwise comparisons between constructs were performed using two-tailed Student’s t-tests with a Tukey HSD adjustment for multiple comparisons. Statistically distinct groups were determined at α = 0.05 and visualized using compact letter displays (CLD)^25^. Groups that share a letter are not statistically different from each other, while groups that do not share any letters are significantly different from each other.

### Western blotting and phosphoprotein staining

Reflectin-transfected cells were washed with PBS and lysed using Thorner buffer (10% glycerol, 8 M urea, 5% (wt/vol) SDS, 40 mM Tris pH 6.8, 4 mg/ml bromophenol blue, supplemented with 5% β-mercaptoethanol)^26^. Lysates were heated at 90°C for 10 minutes, then separated by running on 4-20% Stain-free Tris-glycine SDS-PAGE (Bio-Rad) gel with Tris-glycine running buffer at 200 V for 30 minutes. Total protein load was quantified using stain-free gel settings on a ChemiDoc MP gel imager (BioRad)^27^. Proteins were transferred to a PVDF membrane using 1X transfer buffer (25 mM Tris, 192 mM glycine) at 70 V for 1 hr. Membrane was blocked in 5% milk in 1X TBST (20 mM Tris, 150 mM NaCl, 0.1% Tween-20), followed by incubation overnight at 4°C with anti-GFP primary antibody (GFP Antibody (B-2), 1:1000; Santa Cruz Biotechnology) or anti-mCherry (RPCA-mCherry; 1:5000; Encor Biotechnology) in 5% milk/TBST. Following three 5-minute washes in TBST, membranes were incubated for 30 minutes at room temperature on a rocker with goat anti-mouse horseradish peroxidase (1:10,000; Jackson ImmunoResearch) or goat anti-rabbit horseradish peroxidase (1:10,000; Jackson ImmunoResearch) in 5% milk/TBST. Membranes were then washed twice in TBST for 5 minutes each. Protein bands were visualized using the ECL Western Blotting Substrate (BioRad).

For phosphoprotein detection using Pro-Q Diamond Blot Stain (Thermo Scientific), sample preparation and staining were performed according to the manufacturer’s protocol. Following Pro-Q staining, membranes were re-wetted with 100% methanol and washed three times in PBS on a rocker for 15 min each at room temperature before immunoblotting for mCherry as described above.

## Results

### Reflectin isoforms partition by subcellular compartment and co-condense via localized heterotypic mixing

*D. opalescens* iridocytes have been shown to contain four expressed reflectin isoforms, two of which are classified as canonical (A1, A2), and two as non-canonical (B, C) (Fig. 1A; sequences of all reflectins in this work are shown in Table S1)^3,4^. Canonical reflectin A1 and A2 exhibit a modular block copolymer architecture consisting of alternating conserved Reflectin Motifs (RMs) and cationic linker regions. In contrast, non-canonical reflectin B and C lack the repeating multi-block domain architecture, instead being dominated by cationic, linker-like regions with a single domain (poorly conserved, in the case of reflectin C). Within canonical reflectins, amino acid compositional profiling indicates that tyrosine residues are heavily concentrated within the linker regions, where they likely operate as primary multivalent stickers driving assembly (through π-π and cation-π interactions)^28^ (Fig. 1B; complete analysis of all WT reflectin isoforms shown in Fig. S1). In contrast with canonical reflectins, the linker regions of non-canonical reflectins are less enriched in these aromatic residues.

When WT reflectins were individually transiently overexpressed as either eGFP or mScarlet fluorescent fusion constructs in HeLa cells, all reflectin isoforms formed micron-scale puncta (eGFP-tagged WT reflectins shown in Fig. 1C, mScarlet-tagged WT reflectins shown in Fig. S2), consistent with the formation of biomolecular condensates and previous observations from overexpression in HEK293/HEK293T cell lines^21,22^. Successful expression of WT reflectins (and expression of all constructs described in this work) was confirmed by western blotting against eGFP or mScarlet (Fig. S3). Recovery of expressed reflectin required the use of denaturing Thorner buffer for detection via western blotting, confirming the low solubility and high condensation propensity of reflectins in living cells. Reflectin isoforms showed differences in partitioning between the nuclear and cytoplasmic compartments that were also generally consistent with previously reported observations^21,22^. Overexpressed reflectin A1 primarily localized to the cytoplasm, while A2, B, and C primarily localized to the nucleus. While the exact mechanisms specifying reflectin compartmental localization are unknown, previous work has shown that this partitioning is length-dependent, with longer reflectins predominantly residing in the cytoplasm and shorter ones in the nucleus^22^. However, condensates of most reflectin isoforms could be observed outside of the primary compartments described above.

To assess whether different reflectins colocalized into heterotypic condensates, we co-transfected different reflectin isoforms where one was eGFP-tagged, and the other mScarlet-tagged. Dual-color microscopy revealed that all combinations of reflectin isoforms localized within the same compartment coalesced readily into blended heterotypic assemblies, and that localization of these heterotypic condensates was consistent with single-reflectin expression patterns (Fig. 2). This indicates that compartmentalization is dominant over the attractive intermolecular forces that drive co-condensation^22^.

**Figure 2.**
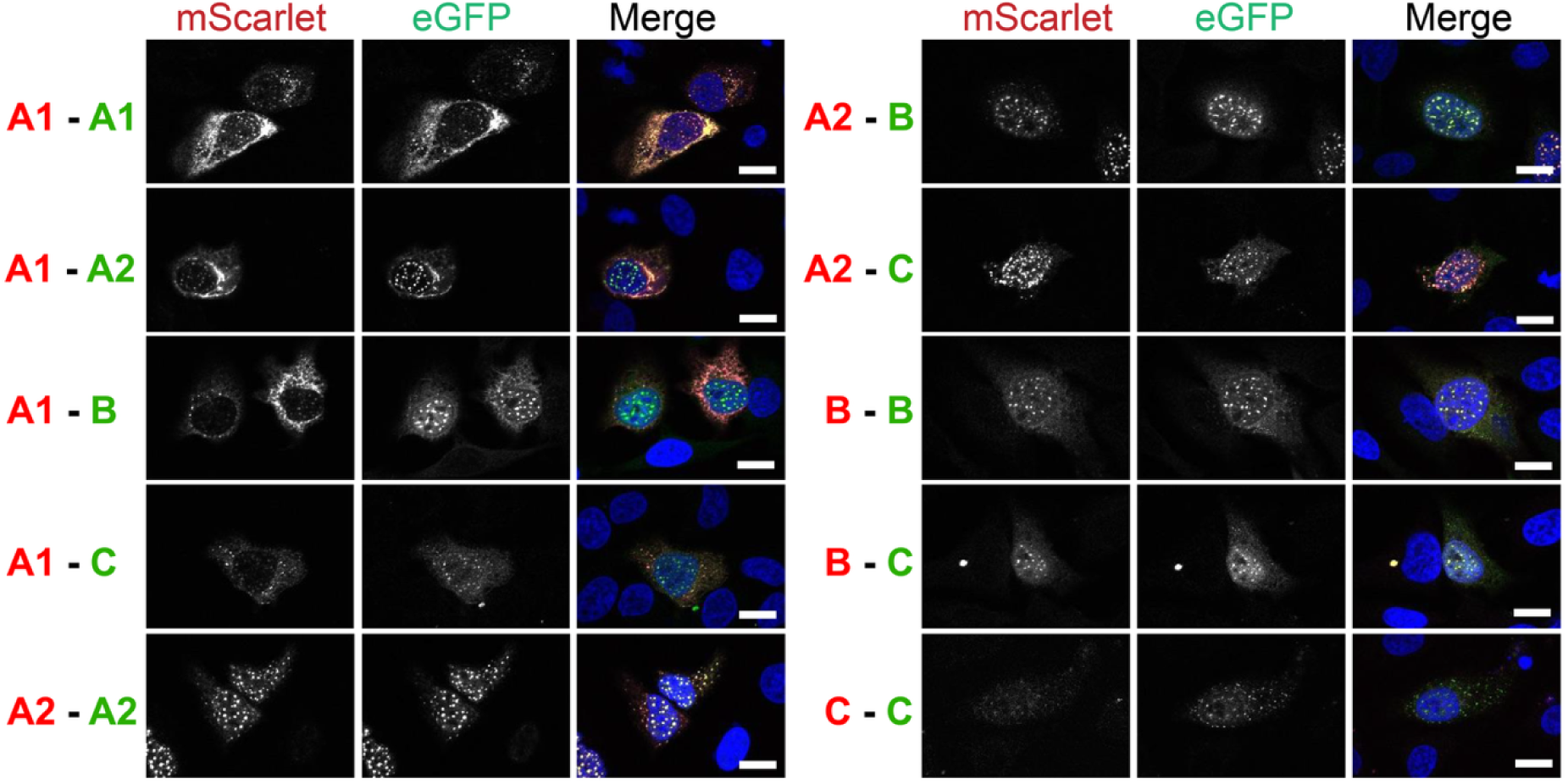
Co-expression images demonstrating co-condensation of localized isoforms. Confocal fixed cell images of eGFP– and mScarlet-reflectin fusions display similar reflectin subtype-dependent localization and assembly morphologies. DAPI nuclear stain is shown as blue in merged images. Scale bar = 20 μm.

Because reflectins were shown to be endogenously phosphorylated in *D. opalescens* iridocytes, we tested for the presence of reflectin phosphorylation when expressed in HeLa cells using a phosphoprotein stain previously used successfully to detect native phosphorylation in squid iridocytes^3^. We were unable to detect phosphorylation of wild-type reflectin isoforms from HeLa cell extracts, suggesting that extensive basal phosphorylation does not occur and that phosphorylation is limited in HeLa, if existent at all (Fig. S4).

### Canonical and non-canonical reflectins demonstrate divergent recovery kinetics

To evaluate the material properties of the observed homotypic reflectin condensates, we performed quantitative whole condensate fluorescence recovery after photobleaching (FRAP) analyses in live cells (Fig. 3). A complete table of fitting data and cell counts for these constructs and other homotypic condensates in this work is provided in Table S2. Non-canonical nuclear reflectin B and C condensates exhibited robust, uniform fluid-like recoveries with relatively minimal Coefficients of Variation (CV) in their mobile fraction (M_f_) (reflectin B M_f_ = 0.85 ± 0.08, CV = 7%; reflectin C M_f_ = 0.85 ± 0.11, CV = 13%) (Fig. 3A-B). Assessment of molecular exchange rates via the recovery half-times (t_1/2_) indicated rapid exchange in these condensates (reflectin B t_1/2_ = 6.74 ± 2.87 s; reflectin C t_1/2_ = 12.29 ± 5.63 s).

**Figure 3.**
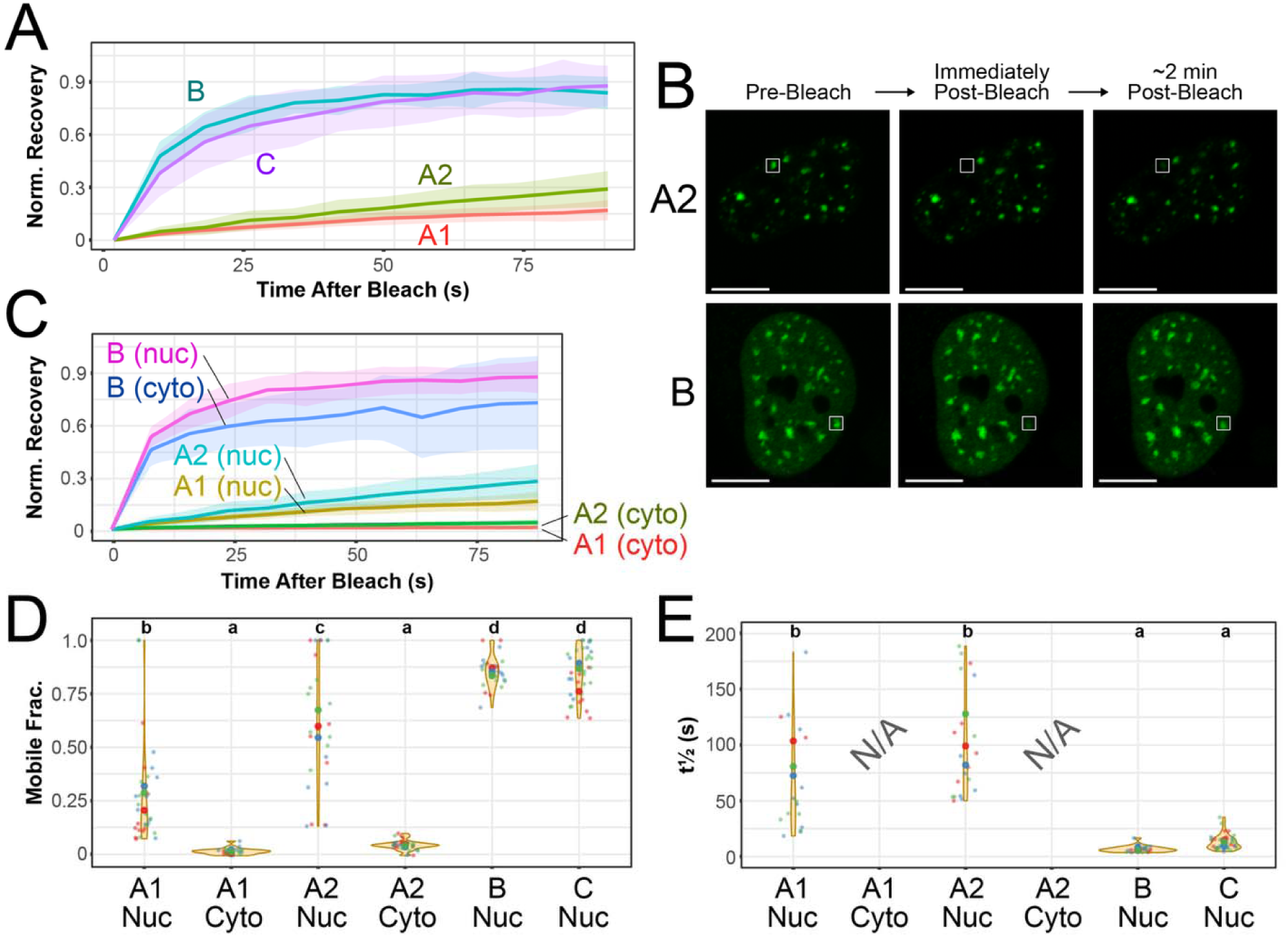
Reflectin isoforms display different dynamics as assessed by FRAP. A) Representative FRAP curves of nuclear condensates of eGFP-tagged WT *D. opalescens* reflectins A1, A2, B, and C. Narrow lines indicate the mean, and shaded areas indicate the standard deviation across time points from a single biological replicate. B) Representative live cell confocal images of a reflectin A2 condensate (top) showing little recovery, and a reflectin B condensate (bottom) showing substantial recovery. Scale bar = 25 μm. Bleached areas are indicated by the white box. C) Representative FRAP curves of A1, A2, and B nuclear or cytoplasmic condensates. A1 and A2 are eGFP-tagged; B is mScarlet-tagged (Fig. S5). Nuclear A1 and A2 curves reproduced from first panel. D) Mobile fraction (M_f_) distributions and E) fluorescence recovery half-time (t_1/2_) distributions after fitting of each individual cell fluorescence recovery curve (*n* = 3 biological replicates). Larger centered dots denote the means of individual biological replicates. Compact letter display (CLD) letters at the top distinguish statistically significant differences between groups within each panel. No t_1/2_ data is shown for A1 [Cyto] and A2 [Cyto], as no condensates had >15% normalized recovery.

In contrast, canonical nuclear reflectin A1 and A2 condensates displayed varied mobile fractions with significant condensate-to-condensate heterogeneity across all biological replicates, with some condensates displaying recovery percentages as low as 10% for A1 and 13% for A2 (Fig. 3A-B). We defined condensates with normalized recovery of <15% as being under dynamic arrest; for these cells, M_f_ was estimated using the final normalized intensity data point. According to this definition of dynamic arrest, 39% of A1 (12/31) and 15% of A2 (4/26) condensates were dynamically arrested (for comparison, 0% of B or C condensates were dynamically arrested). Overall, A1 showed a lower average M_f_ than A2 but similarly high variability (A1 M_f_ = 0.27 ± 0.24, CV = 89%; A2 M_f_ = 0.61 ± 0.30, CV = 49%) (Fig. 3D). Recovery half-times (t_1/2_) for the dynamic components of A1 and A2 were significantly prolonged and variable (A1 t_1/2_ = 80 ± 71 s; A2 t_1/2_ = 106 ± 51 s) relative to non-canonical assemblies discussed above (Fig. 3E). In whole, our FRAP analyses demonstrate that canonical reflectins undergo substantially slower molecular exchange and undergo solidification far more readily than non-canonical reflectins and are likely sensitive to small differences in intracellular conditions that drive solidification at variable rates.

As physiological reflectin lamellae are not localized to the nucleus in cephalopod iridocytes^4,6,29^, we assessed whether condensate localization within the nucleus or cytoplasm significantly impacted dynamics by measuring FRAP curves of cytoplasmic A1, A2, and B condensates (Fig. 3C). Results showed that cytoplasmic condensates had even less recovery than nuclear-localized ones, with almost all A1 and A2 condensates being dynamically arrested according to our <15% recovery definition. In contrast with A1 and A2, cytoplasmic mScarlet-B condensates displayed only a small but significant decrease in M_f_ (but not t_1/2_) relative to nuclear-localized ones (mScarlet-B [nuclear] M_f_ = 0.88 ± 0.08, t_1/2_ = 6.8 ± 2.9 s vs. mScarlet-B [cytoplasmic] M_f_ = 0.74 ± 0.16, t_1/2_ = 7.3 ± 5.13 s; *P_Mf_* < 0.001, *P_t_*_1/2_> 0.999) (Fig. S5), showing that not all cytoplasmic reflectin condensates undergo arrest.

### Dissecting sequence architecture identifies linkers as primary drivers of molecular immobility

To determine whether the dynamic arrest observed in block copolymeric canonical reflectins is encoded by specific sequence blocks, we characterized reflectin A1 truncation variants consisting of either isolated concatenated conserved reflectin motifs (A1 RM) or isolated linker (A1 LO) regions (Fig. 4A). Consistent with previous definitions, we identified the reflectin motif regions as composed of the highly conserved unique N-terminal domain and the canonical domain regions defined by the conserved core M/FD(X)_5_MD(X)_5_MDX_3/4_, extended outwards from this core to conserved prolines that bracket these regions (Table S1)^3,9^. Linkers were defined as the regions interspersed between the conserved motifs (Table S1). Confocal imaging showed that both A1 RM and A1 LO constructs formed nuclear condensates (Fig. 4B). FRAP analysis of the A1 LO construct revealed low condensate mobility, most quantitatively alike to A2 WT (A1 LO M_f_ = 0.31 ± 0.13, t_1/2_ = 53.04 ± 38.49 s) (Fig. 4C). We also tested an additional A1 LO construct that expanded the linker regions outwards an additional 4 amino acids per block (A1 LO-expanded); this construct exhibited essentially the same dynamics by FRAP (Fig. S6), demonstrating that low fluidity is an intrinsic property of linker composition rather than a boundary artifact from specific block assignment. In contrast with the linker-only constructs, an isolated A1 RM construct exhibited significant fluorescence recovery, indicating far greater fluidity in these condensates (A1 RM M_f_ = 0.92 ± 0.07, t_1/2_ = 4.78 ± 1.92 s) (Fig. 4C). Lastly, we characterized a construct containing the cationic N-terminal linker of A1 together with all domains (A1 LnRM, Fig. 4A); this construct displayed a slight but statistically insignificant decrease in M_f_ (*P_Mf_* = 0.134, *P_t_*_1/2_ = 0.913) relative to A1 RM (Fig. 4F-G). In summary, these results suggest that the linker regions interspersed between the conserved reflectin domains are the primary factor driving slow dynamics and dynamic arrest within canonical reflectin condensates, consistent with the enrichment in these regions of arginine and tyrosine residues (Fig. 1B).

**Figure 4.**
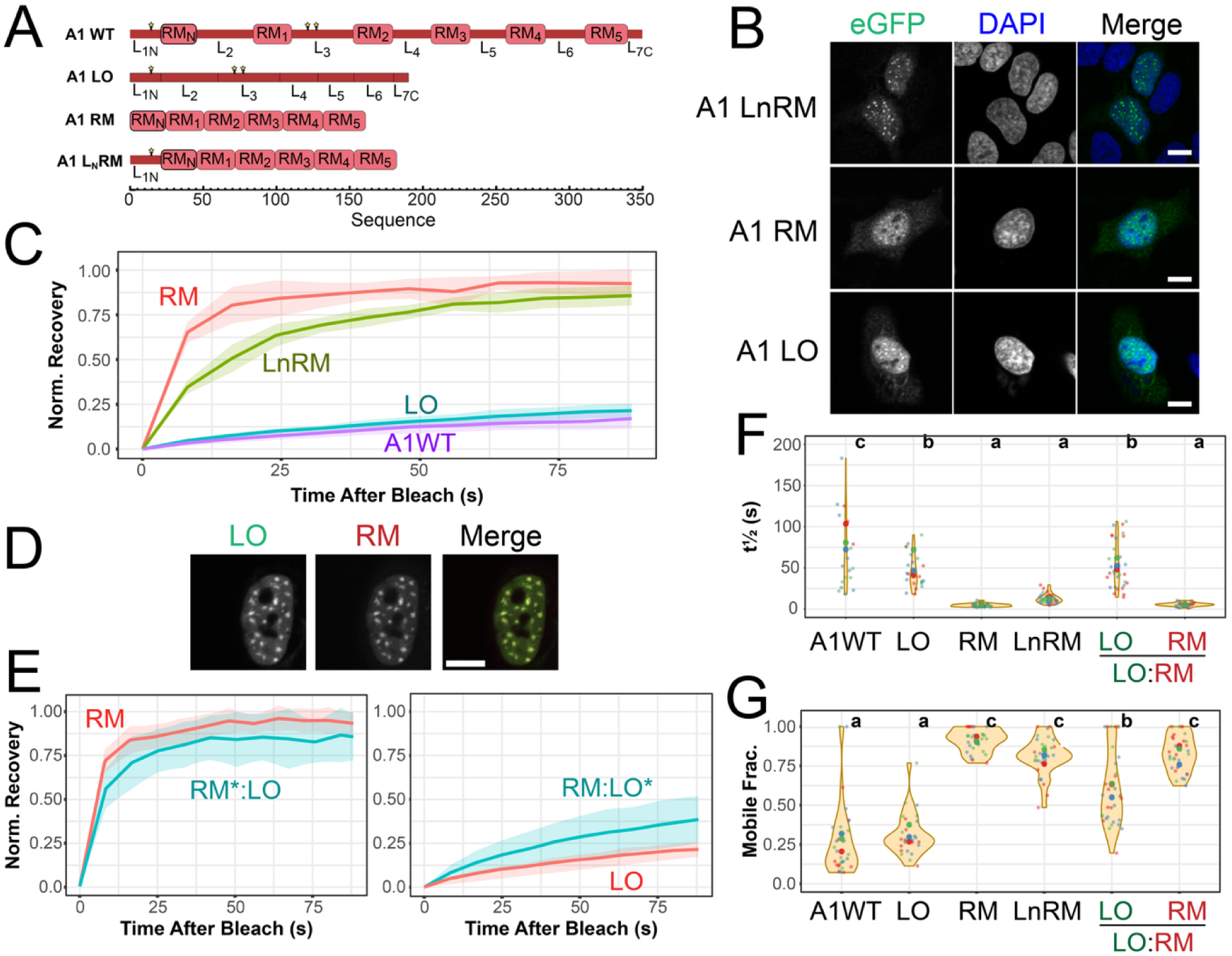
Linker and reflectin motif-only blocks contribute unequally to the material properties of canonical reflectins. A) Schematics of single or mixed block mutants analyzed in this work, compared to A1 WT. B) Fixed cell confocal images of eGFP-tagged A1 block mutants. Scale bar = 10 μm. C) Representative FRAP curves of A1 block mutants, compared to nuclear-localized A1 WT. D) Live cell confocal images of co-transfected A1 RM and A1 LO constructs, demonstrating co-localization. Scale bar = 10 μm. E) Representative FRAP curves of A1 RM and A1 LO constructs within heterotypic condensates. Single-component controls for comparison reproduced from panel C. F) M_f_ and G) t_1/2_ distributions after fitting of each individual cell fluorescence recovery curve (*n* = 3 biological replicates). Larger centered dots denote the means of individual biological replicates. CLD letters at the top distinguish statistically significant differences between groups within each individual panel. A1WT data reproduced from Fig. 3 for comparison.

To assess whether A1 RM and A1 LO constructs form co-condensates, we co-transfected HeLa cells with both constructs tagged to different fluorescent proteins. A1 LO and RM constructs colocalized into heterotypic LO:RM condensates, demonstrating that the overall physicochemical sequence grammar shared by the blocks can drive co-recruitment (Fig. 4D). FRAP analysis of these dual-component condensates reveals mutual buffering of fluidity (Fig. 4E), yielding a slight but statistically significant increase in M_f_ of the LO*:RM component (LO*:RM M_f_ = 0.60 ± 0.23 vs. 0.31 ± 0.13, *P_Mf_* < 0.001, *P_t_*_1/2_ > 0.999), alongside a slight, non-significant reduction in the M_f_ of the more fluid LO:RM* component (LO:RM* M_f_ = 0.83 ± 0.11 vs. RM 0.92 ± 0.07, *P_Mf_* = 0.364, *P_t_*_1/2_ > 0.999) relative to the isolated constructs (Fig. 4F-G, complete data set for all heterotypic condensates in Table S3).

### Phosphomimetic mutations progressively fluidize some reflectin condensates

ACh signaling in tunable iridocytes triggers phosphorylation of canonical reflectins A1 and A2, and dephosphorylation of noncanonical reflectin B^3,4,15^. Previous *in vitro* work with purified reflectins shows that phosphomimetic polyglutamate insertions led to assembly of monomeric protein into spherical, dynamically arrested assemblies of a size calibrated to the overall degree of protein net neutralization^17^. To assess the effects of phosphorylation on reflectin condensates, we generated a series of phosphomimetic mutants of reflectins A1, A2, and B with increasing numbers of introduced glutamates in linear series (Y or S E (1E), EEE (3E), or EEEEE (5E)). The sites chosen for these phosphomimetic mutations were previously identified physiological phosphorylation sites, with four for A2 (Y12/Y214/S218/Y223), and two for A1 (Y14/Y128) and B (S228/S259) (Fig. 1A). While two physiological sites have been identified for reflectin B and three sites for A1^3,4^, only two sites in A1 were used here to replicate the previous *in vitro* experiments^17^.

All phosphomimetic mutants formed condensates with similar localization and general morphology as WT proteins (Fig. S7). As assessed by FRAP, incremental addition of negative charges via mutagenesis transformed heterogenous reflectin A2 condensates into a more homogeneous, progressively more fluid state (Fig. 5A). A2 3E and 5E mutants showed a statistically significant increase in M_f_ relative to A2 WT and 1E (A2 5E M_f_ = 0.80 ± 0.10, CV = 13%, p < 0.005) and had zero individual dynamically arrested condensates according to the <15% recovery threshold (Fig. 5A, Table S2). Additionally, a progressive decrease in t_1/2_ was observed across the phosphomimetic series (A2 WT t_1/2_ = 106.2 ± 51.38 s; A2 1E t_1/2_ = 78.94 ± 50.46 s; A2 3E t_1/2_ = 46.01 ± 32.25 s; A2 5E t_1/2_ = 14.88 ± 5.74 s), indicating increasing molecular exchange within the mobile fraction (Table S2).

**Figure 5.**
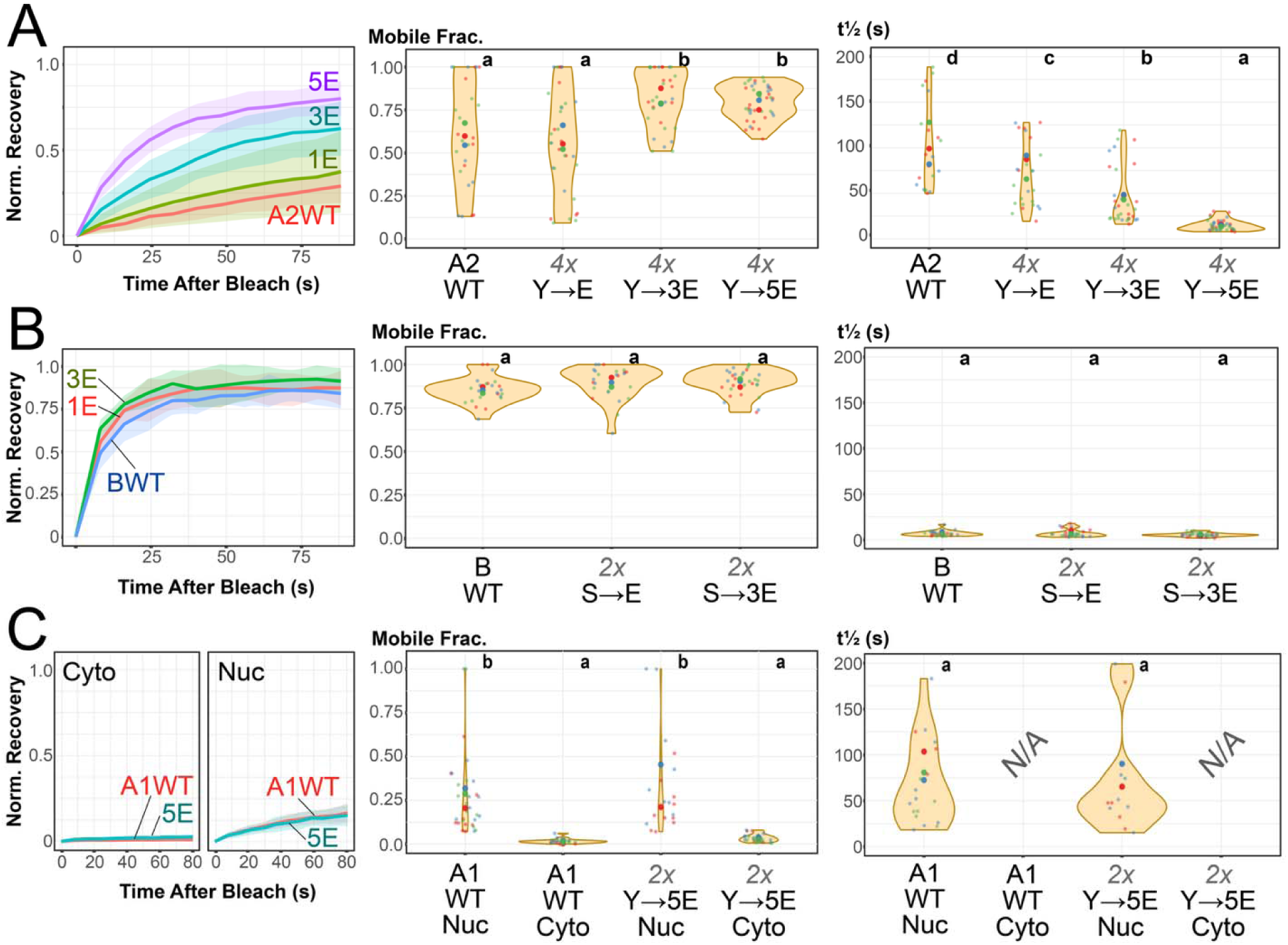
In general, phosphomimetic reflectin mutants display incrementally increased internal dynamics. A) Representative FRAP curves and fitted parameters of A) Reflectin A2 (nuclear), B) Reflectin B (nuclear), and C) Reflectin A1 (nuclear and cytoplasmic) WT proteins and their phosphomimetic mutants. Leftmost panels for each protein show representative FRAP curves; middle and right panels show M_f_ and t_1/2_ distributions after fitting of each individual cell fluorescence recovery curve (*n* = 3 biological replicates). Larger centered dots denote the means of individual biological replicates. CLD letters at the top distinguish statistically significant differences between groups within each panel. A1 WT, A2 WT, and B WT data reproduced from Fig. 3 for comparison.

In contrast with A2, reflectin B exhibited a slight, though statistically insignificant, increase from B WT in M_f_ (B 3E M_f_ = 0.93 ± 0.06, p = 0.099) and decrease in t_1/2_ (B 3E t_1/2_ = 5.29 ± 1.89 s, p = 0.197) across the phosphomimetic series compared to the wild-type baseline (Fig. 5B, Table S2). This lack of significant effect is likely attributable to the preexisting greater fluidity of the WT reflectin B protein. Lastly, we tested the effects of phosphomimetic mutations on both nuclear and cytoplasmic reflectin A1. A1 5E phosphomimetic condensates in either compartment did not display observable increases in recovery compared to homotypic A1 WT condensates in the same compartment (Fig. 5C). The lack of responsiveness of A1 to these phosphomimetic mutations may be due to the increased length and valency of A1 compared to A2, an insufficient number of introduced phosphomimetic sites to overcome interactions driving solidification in the homotypic A1 condensates, or other sequence differences between the canonical proteins.

### Heterotypic reflectin isoform mixing disrupts homotypic crosslinks to increase condensate fluidity

In tunable cephalopod iridocytes, canonical and non-canonical reflectins co-exist within the same lamellar structures^4^. To examine potential heterotypic effects from mixing reflectins, we measured FRAP recovery curves of co-condensates formed by co-expressing diverse combinations of canonical and noncanonical reflectin isoforms. These combinations included both two canonicals together (A1:A2), as well as multiple combinations of canonical and noncanonical reflectins.

Heterotypic nuclear condensates formed from eGFP-tagged reflectin A1 together with mScarlet-tagged reflectin A2 did not show an increase in fluidity relative to their independent homotypic condensates, with most cells being dynamically arrested (20/31 for A1, 26/31 for A2) as defined by normalized recovery < 15% (Fig. S8, Table S3). In contrast, heterotypic condensates composed of reflectin A2 mixed with B or C showed substantial though heterogenous increases in A2 fluidity compared to A2 homotypic condensates, displaying statistically significant decreases in t_1/2_ of the mobile fraction relative to A2 WT (A2*:B t_1/2_ = 51.23 ± 42.22 s, p < 0.001; A2*:C t_1/2_ = 71.10 ± 35.46 s, p = 0.020), and statistically insignificant increases in M_f_ (A2*:B M_f_ = 0.68 ± 0.27, p = 0.568; A2*:C M_f_ = 0.78 ± 0.21, p = 0.074) (Fig. 6A,C-D, Table S3). These results indicate that non-canonical reflectin B or C incorporation leads to increases in fluidity of the dynamic canonical reflectin component within the heterotypic condensates. In contrast with the A2 component, neither reflectin B nor C showed significant changes in their recovery curves within the A2 heterotypic condensates compared to their homotypic condensates (Fig. 6B, Fig. S9).

**Figure 6.**
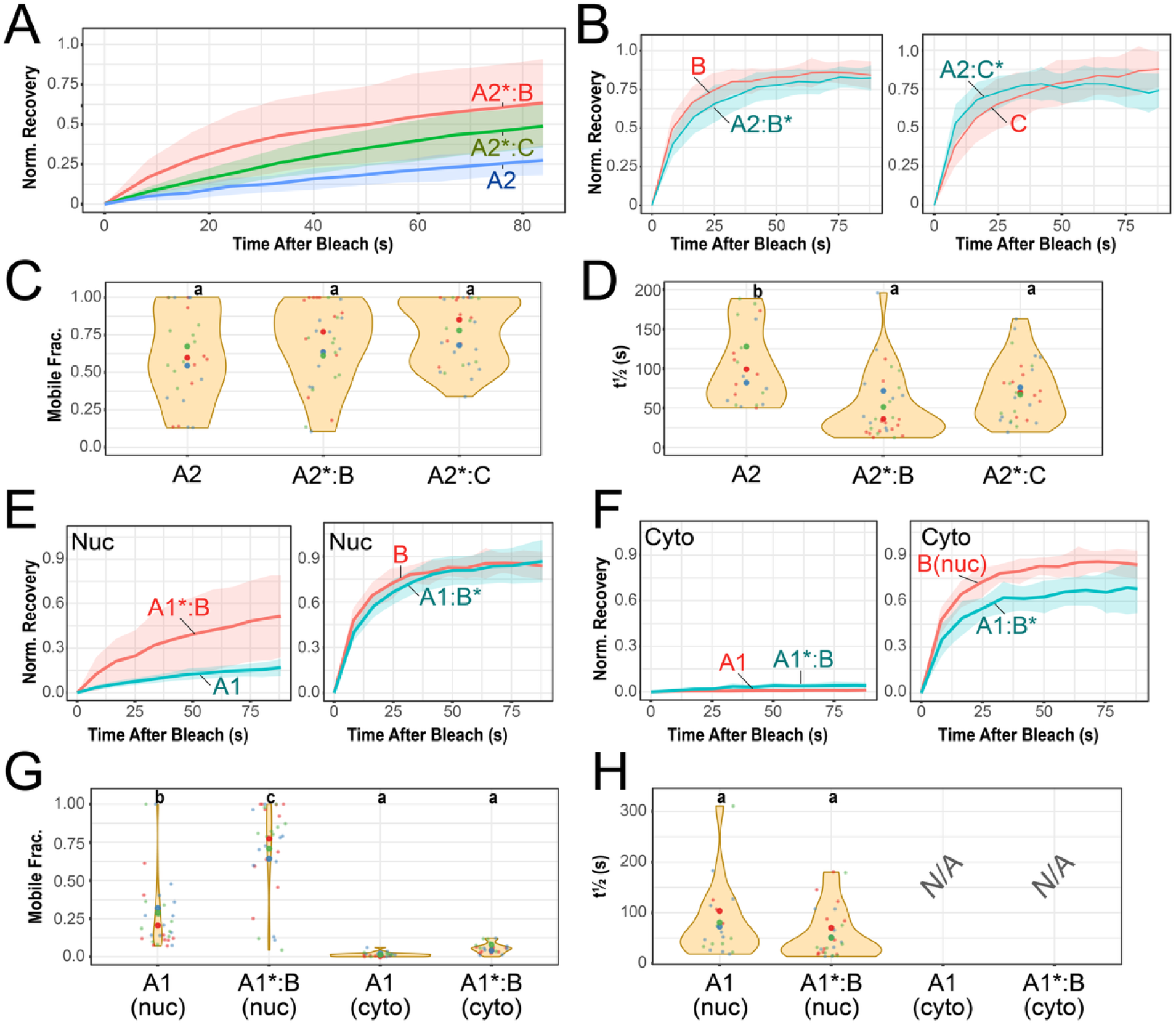
Incorporation of noncanonical reflectins modulates heterotypic condensate fluidity. A) FRAP curves of components within a heterotypic condensate of A2 (left panel) with either B or C, compared to A2 alone (reproduced from Fig. 3). B) FRAP curves of B or C components within heterotypic condensates with A2, compared with their homotypic condensates (reproduced from Fig. 3) C) M_f_ and D) t_1/2_ distributions after fitting of A2 component within heterotypic condensates with B or C after fitting of each individual cell fluorescence recovery curve (*n* = 3 biological replicates). Larger centered dots denote the means of individual biological replicates. CLD letters at the top distinguish statistically significant differences between groups within each panel. E) FRAP curves of individual components of A1:B heterotypic condensates in the nucleus or the F) cytoplasm. G) M_f_ and H) t_1/2_ distributions after fitting of A1 component in homotypic vs. heterotypic A1:B condensates across nuclear or cytoplasmic compartments (*n* = 3 biological replicates). Homotypic A1, A2, and B data reproduced from Fig. 3 for comparison. No t_1/2_ data is shown for A1 [Cyto] and A1*:B [Cyto], as no condensates had >15% normalized recovery.

Heterotypic nuclear condensates of reflectin A1 and B also showed a significant increase in M_f_, but not t_1/2,_ of the A1 component compared to homotypic nuclear A1 condensates (A1*:B M_f_ = 0.71 ± 0.30, p < 0.001; A1*:B t_1/2_ = 58.76 ± 51.05 s, p = 0.267) with the B-mScarlet component showing a 38.3% increase in t_1/2_ from B-mScarlet alone (*P_t_*_1/2_ = 0.001, *P_Mf_* = 1.0), which may be due to a viscous drag effect from the A1 component (Fig. 6E, Fig. S9, Table S3). In contrast, cytoplasmic A1:B condensates did not show an increase in fluorescence recovery of the A1 component. Importantly, despite the complete lack of recovery in the A1 component of these heterotypic condensates, the B component of these cytoplasmic heterotypic condensates did demonstrate fluorescence recovery, similar to cytoplasmic B-mScarlet condensates by themselves (Fig. 6F, Fig. S9). This observed fluorescence recovery of the B component within these cytoplasmic heterotypic condensates suggests that the lack of canonical reflectin recovery observed within cytoplasmic condensates may be driven by their low inter-condensate reflectin exchange rather than total dynamic arrest within these condensates.

Lastly, we sought to create two-component heterotypic condensates composed of canonical A2 and noncanonical B that mimicked the native phosphorylated and unphosphorylated reflectin composition found in activated and unactivated iridocytes^4^ (Fig. 7). To mimic the noniridescent, unactivated lamellar environment of a tunable iridocyte, we co-transfected A2 WT and the B-3E phosphomimetic constructs. To mimic the iridescent, activated lamellar environment, we co-transfected A2-3E phosphomimetic and B WT. Physiomimetic heterotypic nuclear condensates resembling activated iridocytes displayed significantly greater reflectin A2 fluidity (A2-3E*:B M_f_ = 0.86 ± 0.14; t_1/2_ = 25.39 ± 14.37 s) than those resembling the unactivated ground state (A2*:B-3E M_f_ = 0.72 ± 0.23, p = 0.049; t_1/2_ = 71.68 ± 58.66 s, p < 0.001) or of A2-3E condensates alone (Table S3), demonstrating that canonical phosphorylation and non-canonical dephosphorylation act synergistically to fluidize these reflectin condensates.

**Figure 7.**
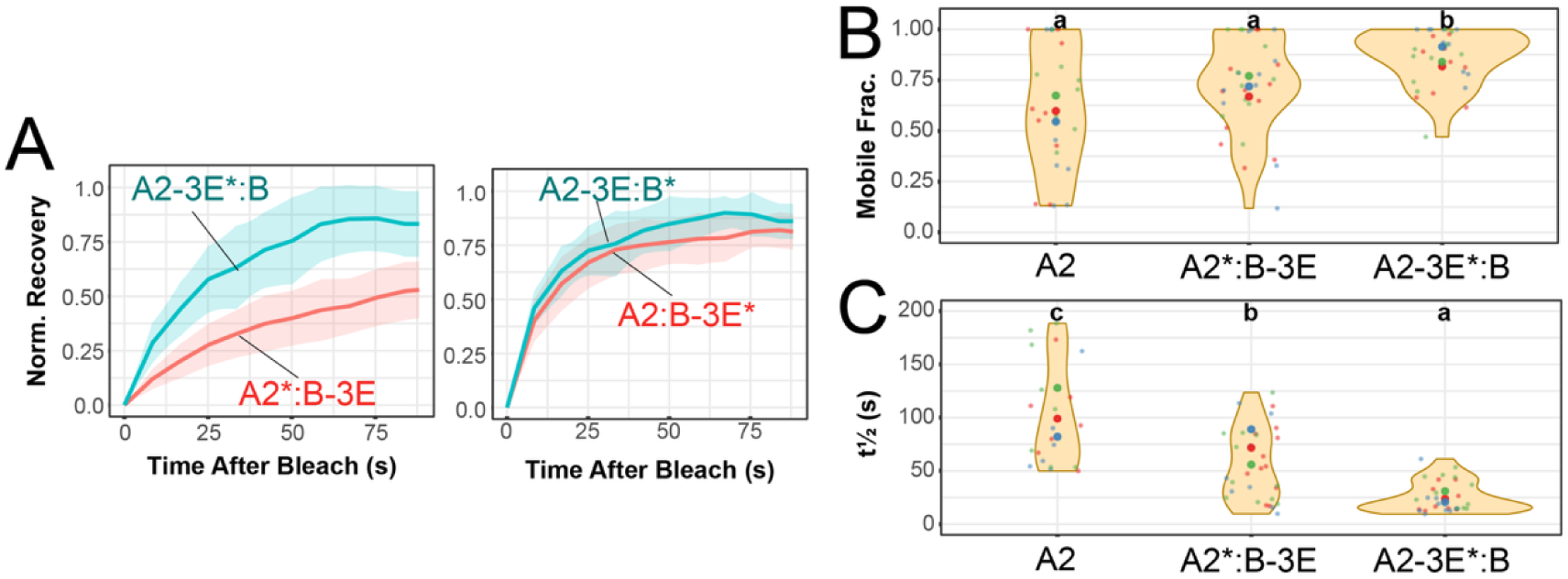
Physiological mixtures of phosphomimetic mutants modulate heterotypic condensate fluidity. A) FRAP curves of physiomimetic heterotypic condensates mimicking the noniridescent, unactivated state (unphosphorylated A2 WT, phosphorylated B-3E) and the iridescent, activated state (phosphorylated A2-3E, unphosphorylated B WT). An asterisk denotes the protein being tracked in that panel. B) M_f_ and C) t_1/2_ distributions after fitting of the A2 component within heterotypic condensates after fitting of each individual cell fluorescence recovery curve (*n* = 3 biological replicates). Larger centered dots denote the means of individual biological replicates. CLD letters at the top distinguish statistically significant differences between groups within each panel. Homotypic A2 data reproduced from Fig. 3 for comparison.

## Discussion

Reflectins assemble within cephalopods to form biophotonic platelets characterized by high protein density and concomitant high refractive index^4,6,30,31^. Studies of tunable iridescence in the squid *Doryteuthis opalescens* established that neuronal acetylcholine (ACh) signaling drives the phosphorylation of canonical A-type reflectins alongside simultaneous dephosphorylation of non-canonical reflectin B^3,5^. Neutralization of net positive charge triggers reflectin condensation within intracellular lamellae, driving water efflux into the extracellular space, with the resulting increase in lamellar refractive index and optical contrast generating dynamic structural coloration within the cellular Bragg reflector^3–5^. This net charge-neutralization mechanism has been extensively modeled *in vitro* with purified proteins, where pH titration of purified monomeric reflectins serves as a surrogate for *in vivo* phosphorylation^16,17,20^. Recombinant reflectin thin films undergo hierarchical assembly that is highly sensitive to environmental cues^10,32–34^, and under low ionic strength, reflectins assemble into dynamically arrested nanoparticles whose dimensions scale with the degree of charge neutralization^17,20,35^. Electrochemical reduction of imidazolium groups in solubilized reflectins and film constructs has further validated the coulombic switch framework^36–38^. However, despite extensive characterization of purified reflectins, the degree to which these *in vitro* phase behaviors describe reflectin assemblies within living cells remains unresolved.

Biomolecular condensates are dynamic liquid or gel assemblies whose composition and material properties are strictly regulated to execute specific cellular functions^39,40^. Aberrant material transitions of condensates have been frequently linked to pathological aggregation^41–43^. Previous work demonstrated that purified reflectins form single– or multi-component micron-sized condensates *in vitro* under high ionic strength with pH-dependent dynamics^17–20^. In contrast to other condensate systems, reflectin execute a unique and specialized biological function: they directly manipulate light by establishing high local refractive indices when packed into higher-order Bragg lamellae^2^. Ultrastructural TEM analysis of tunable squid iridophores has previously revealed that reflectin platelets undergo a functional state change from a gel-like resting phase to a fluid-like (sol) activated phase, suggesting that the optical output of the Bragg lamellae is coupled to alterations in lamellar material state^29^. By evaluating wild-type and mutant reflectin condensates within a crowded cellular environment, our work bridges a gap between simplified *in vitro* thermodynamics and intracellular biophysics.

Our findings uncover sequence features that govern dynamics of cellular reflectin condensates. Quantitative whole condensate FRAP characterization of single-component wild-type reflectin condensates reveals striking dynamic divergence across isoforms: canonical A-type reflectins display a low molecular exchange, dynamically arrested state, whereas non-canonical variants exhibit rapid exchange. Canonical A-type reflectins feature a block-copolymeric sequence architecture consisting of ampholytic, evolutionarily well-conserved domains (Reflectin Motifs, RMs) interspersed with cationic, tyrosine-rich linkers. Direct comparison of FRAP curves for reflectin motif-only (A1 RM) and linker-only (A1 LO) constructs indicates that the physicochemical drivers of low exchange reside primarily within the linker regions of the reflectins. These observed differences between A1 LO and A1 RM constructs result from compositional differences between the two regions: the linkers are heavily enriched in tyrosine and arginine residues, which function as primary multivalent stickers (engaging in π-π and cation-π interactions) that drive low molecular exchange and dynamic arrest^28,44,45^. Notably, co-expression of A1 RM and A1 LO still results in complete co-localization within shared condensates, demonstrating that their shared molecular grammar is sufficient to drive heterotypic co-condensation. These results suggest that the primary functional role of the highly enriched tyrosine-rich linkers is to supply a high density of multivalent crosslinks that promote network percolation and solidification, whereas the conserved reflectin motifs complementarily contribute to the broad phase-separation propensity of these proteins and likely serve an additional physiological role yet to be clearly demonstrated^9,46,47^. Despite being depleted in conserved reflectin motifs and enriched in “linker” regions, non-canonical reflectins B and C display fluorescence recovery kinetics more akin to the conserved motifs of canonical A-type reflectins than their linkers. This disparity stems from the marked compositional differences between canonical and non-canonical linker regions. For example, linkers in canonical A-type reflectins are heavily tyrosine-enriched compared to those of non-canonical variants (*e.g.*, A1 LO 16.5% Tyr vs. B WT 5.8% Tyr). This relative depletion of sticker residues in noncanonical reflectins correlates well with the greater lamellar fluidity previously observed in extracts from activated tunable iridocytes, which are enriched in noncanonical reflectins B and C^4,29^.

Our results demonstrate that phosphorylation modulates condensate dynamics in both homotypic and heterotypic cellular environments. Introducing glutamate residues at four physiological phosphorylation sites drove a progressive increase in molecular exchange within reflectin A2 condensates, aligning with the emergence of iridescence in tunable cells. Previous work has shown that phosphomimetic mutations can tune the size of dynamically arrested reflectin assemblies produced *in vitro* from monomeric reflectin via pH neutralization as a surrogate for phosphorylation, and modeling has shown that the assembly size can be captured through a short-range attraction, long-range repulsion (SA-LR) model^35^. In contrast, introducing phosphomimetic substitutions into homotypic reflectin B and A1 condensates did not yield significant shifts in FRAP kinetics, though the underlying molecular explanations for each protein are likely distinct. While wild-type reflectin B already exhibits high baseline fluidity with less room for further fluidization, wild-type reflectin A1 maintains a largely arrested, low-exchange state. The lack of responsiveness in A1 likely reflects its substantially higher chain valency and sequence length (350 residues in A1 and 5 canonical motifs vs. 230 residues and 3 motifs in A2), which makes introducing two phosphomimetic sites insufficient to overcome the high density of unbuffered linker stickers. In unactivated, noniridescent iridocytes, differences in fluidity between A1 and noncanonical reflectins (including phospho-B) may drive the formation of the irregular, particulate gel structure observed in the ground state of those cells (Fig. 8).

**Figure 8.**
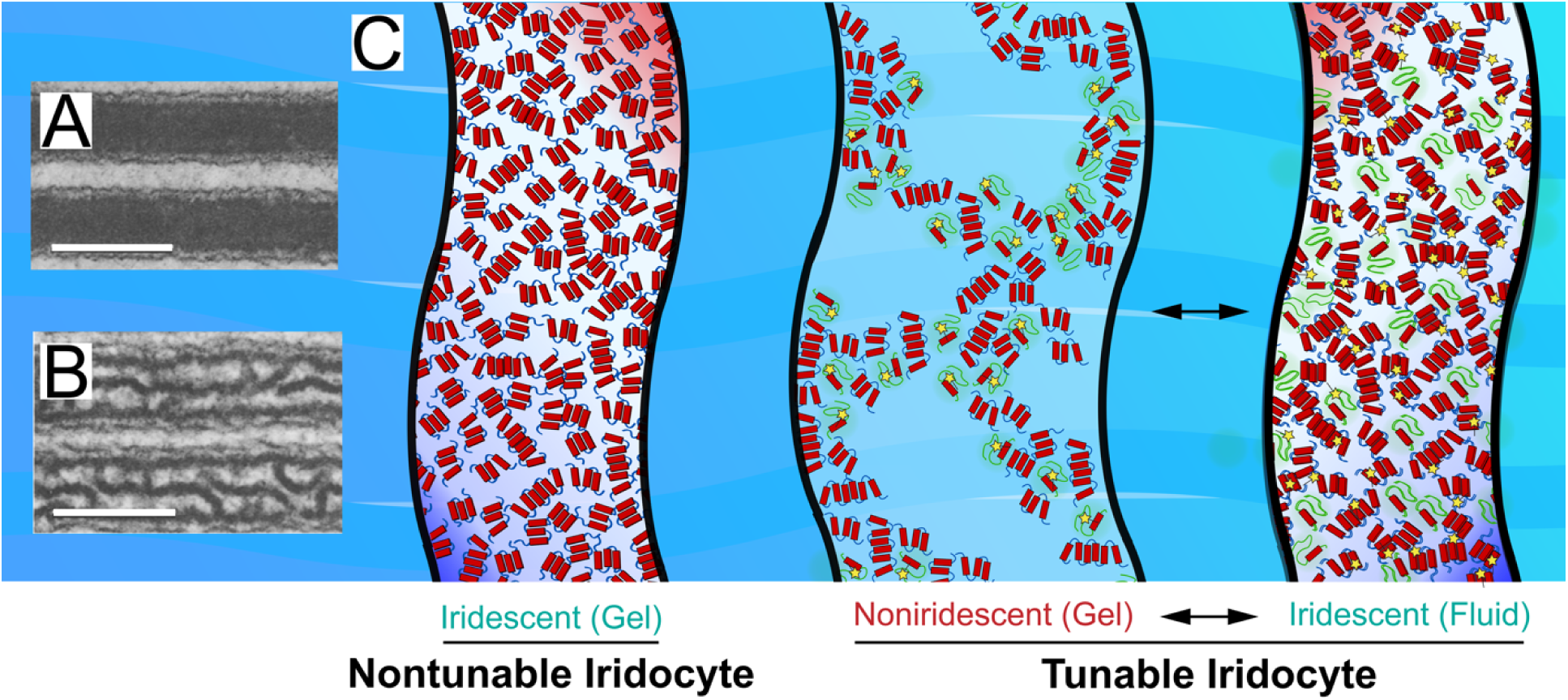
Models of reflectin lamellae in nontunable and tunable iridocytes. Electron micrographs of fixed, tunable iridocytes in an A) ACh-activated state or B) inactive state. Scale bar = 0.25 μm. Panels A and B are reproduced with permission from Springer Media from Cooper *et al.*, Physiological color change in squid iridophores, *Cell. Tiss. Res.*, 1990, 259:15-24^29^. C) Models of reflectin-filled lamellae in nontunable and tunable iridocytes. Canonical reflectins are represented with blue backbones; green backbones represent noncanonical reflectins. Phosphorylation sites represented with yellow stars. Material states of lamellae are designated in parentheses^5,29,53^.

Intriguingly, canonical reflectin A1 and A2 nuclear-localized counterparts exhibit measurably higher fluidity than cytoplasmic condensates. As whole-condensate photobleaching FRAP cannot fully distinguish intra-condensate molecular exchange and inter-condensate exchange via the dilute phase, the lack of recovery of cytoplasmic A1 and A2 may reflect either slow intra-condensate diffusion or a negligibly low dilute-phase concentration (c_sat_)^13^. While definitively isolating intra-from inter-condensate dynamics will require future investigation, in the absence of observable microphase separation the rapid mobility of reflectin B within cytoplasmic co-condensates argues against complete dynamic arrest and loss of intracondensate diffusion, instead suggesting that suppressed inter-condensate exchange underlies the apparent dynamic arrest of cytoplasmic canonical reflectins. The nuclear environment, enriched in polyanionic DNA, nuclear RNAs, and high concentrations of free ATP acting as a biological hydrotrope, may act to increase c_sat_ and plasticize these condensates^48–50^. By forming competitive electrostatic interactions with cationic reflectins, these nuclear polyanions may disrupt homotypic crosslinks and suppress complete solidification. Analogous molecular plasticizers may operate within specialized, non-nuclear Bragg lamellae of cephalopod iridocytes to maintain baseline fluidity required for regulated lamellar assembly.

Heterotypic reflectin interactions represent another critical regulatory tier. Canonical reflectin condensates demonstrate low molecular exchange by FRAP, consistent with the formation of static platelets that assemble into Bragg reflector (or sinusoidal) structures within static iridocytes^6,29^. We observed that reflectin isoforms readily co-condense to form mixed assemblies, and that co-condensation with reflectin B or C significantly enhanced exchange of the A2 component within the co-condensate. Furthermore, two-component physiomimetics of unactivated and activated lamellar states demonstrated clear state-dependent shifts in reflectin A2 exchange. While these simplified, heterologous two-component mixtures do not fully recapitulate the formation of the gel phase observed in native iridocyte lamellae, they correlate well with physiological observations and support the hypothesis that tunable iridescence is directly coupled to transitions in lamellar material state. The amplified change in material properties observed in native iridocytes likely reflects additional physiological factors, including the mixed four-isoform composition which may drive microphase separation^51^, chemical distinctions between native phosphorylation and phosphomimetic surrogates, physical confinement within dense lamellar membranes, and specialized cellular regulatory machinery.

The mechanisms by which phosphorylation and non-canonical isoform inclusion dictate condensate material properties can be unified within a stickers-and-spacers framework grounded in polymer physics ^12,13^. Our findings suggest that the emergence of iridescence in tunable iridocytes represents a fundamental material-state transition, specifically a reversible shift from a low-fluidity particulate gel to a dynamic, highly mobile fluid phase. In this framework, unphosphorylated canonical reflectins form high-valency, heavily crosslinked networks that undergo dynamic arrest into percolated gels with low or no fluidity ^13,52^. Conversely, non-canonical reflectins possess lower sticker valencies and thus act as plasticizers, competitively disrupting contiguous homotypic crosslinks to reduce effective network crosslink density to enhance fluidity^13,49,52^. Within this porous matrix, reversible phosphorylation modulates effective crosslink density to fluidize the assembly, enabling dynamic, reversible switching between a low-fluidity gel and an optically active condensed state coupled to water efflux^5^. While the presence of the noncanonical reflectins within iridocytes is insufficient to drive fluidization in the unactivated state, yielding instead a particulate lamellar gel, upon ACh-activation and subsequent phosphorylation and dephosphorylation, changes in the distribution of attractive and repulsive interactions are sufficient to drive the formation of a homogeneous dense fluid of greater refractive index compared to the extracellular medium^29^ (Fig. 8). In whole, this model provides a straightforward biophysical explanation for how neural signaling triggers material transitions, enabling dynamic camouflage and intraspecies communication in squid.

## Data Acknowledgement

Data are available to be shared upon request. Please contact the corresponding author at.

## Acknowledgement*s*

This material is based upon work supported by the U.S. National Science Foundation under Grant No. 2233670 (to R.L.). Additional support was provided by the John Stauffer Charitable Trust and Soka University.

