## Supplemental Materials for "Heterotypic interactions and sequence features modulate cellular reflectin condensate dynamics"

**Phan, Chi[1]; Watanabe, Rika[1]; Le, Vinh[1]; Walsh, Susan[1]; and Robert Levenson[1]***

**[1]Life Sciences Concentration, Soka University of America, Aliso Viejo, CA 92656, USA**

**Supplemental Table 1. Sequences of WT and mutant reflectins used in this work.** UniProt accession numbers and links for WT reflectins are provided in the table.

**Legend:**

**Linkers: Yellow and Green**

**Reflectin Motifs: Cyan & Magenta**

**GMXX (Reflectin C only): Gray**

**Physiological Phosphorylation & Phosphomimetic Mutagenesis Sites: Red, Underlined**

| Reflectin | Sequence |
|---|---|
| **A1 WT**<br><br>UniProt Accession:<br>A0A088MEP7 | MNRYLNRQRLYNM**Y**RNKYRGVMEPMSRMTMDFQGRYMDSQGRMVDPRYYDHYGRMHDY<br>DRYYGRSMFNQGHSMDSQRYGGWMDNPERYMDMSGYQMDMQGRWMDAQGRYNNPFSQM<br>WHSRQGHYPG**Y**MSHHSMYGRNMHYPYHSHSASRHFDSPERWMDMSGYQMDMQGRWMDN<br>YGRYVNPFHHHMYGRNMFYPYGSHCNNRHMEHPERYMDMSGYQMDMQGRWMDTHGRHC<br>NPLGQMWHNRHGYYPGHPHGRNMFQPERWMDMSSYQMDMQGRWMDNYGRYVNPFSHNY<br>GRHMNYPGGHYNYHHGRYMNHPERQMDMSGYQMDMHGRWMDNQGRYIDNFDRNYYDYH<br>MY* |
| **A1 ReflectinMotifOnly** | MEPMSRMTMDFQGRYMDSQGRMVDPPERYMDMSGYQMDMQGRWMDAQGRYNNPPERWM<br>DMSGYQMDMQGRWMDNYGRYVNPPERYMDMSGYQMDMQGRWMDTHGRHCNPPERWMDM<br>SSYQMDMQGRWMDNYGRYVNPPERQMDMSGYQMDMHGRWMDNQGRYIDNF* |
| **A1 LinkerOnly** | MNRYLNRQRLYNMYRNKYRGVRYYDHYGRMHDYDRYYGRSMFNQGHSMDSQRYGGWMD<br>NFSQMWHSRQGHYPGYMSHHSMYGRNMHYPYHSHSASRHFDSFHHHMYGRNMFYPYGS<br>HCNNRHMEHLGQMWHNRHGYYPGHPHGRNMFQFSHNYGRHMNYPGGHYNYHHGRYMNH<br>DRNYYDYHMY* |
| **A1 LinkerOnly(Expanded)**<br>Added segments:<br>**magenta** / **cyan** | MNRYLNRQRLYNMYRNKYRGVRYYDHYGRMHDYDRYYGRSMFNQGHSMDSQRYGGWMD<br>N**YNNP**FSQMWHSRQGHYPGYMSHHSMYGRNMHYPYHSHSASRHFDS**YVNP**FHHHMYGR<br>NMFYPYGSHCNNRHMEH**HCNP**LGQMWHNRHGYYPGHPHGRNMFQ**YVNP**FSHNYGRHMN<br>YPGGHYNYHHGRYMNH**YIDNF**DRNYYDYHMY* |
| **A1 LnDomainOnly** | MNRYLNRQRLYNMYRNKYRGVMEPMSRMTMDFQGRYMDSQGRMVDPPERYMDMSGYQM<br>DMQGRWMDAQGRYNNPPERWMDMSGYQMDMQGRWMDNYGRYVNPPERYMDMSGYQMDM<br>QGRWMDTHGRHCNPPERWMDMSSYQMDMQGRWMDNYGRYVNPPERQMDMSGYQMDMHG<br>RWMDNQGRYIDNF* |
| **A1 Y2EEE** | MNRYLNRQRLYNM**EEE**RNKYRGVMEPMSRMTMDFQGRYMDSQGRMVDPRYYDHYGRMH<br>DYDRYYGRSMFNQGHSMDSQRYGGWMDNPERYMDMSGYQMDMQGRWMDAQGRYNNPFS<br>QMWHSRQGHYPG**EEE**MSHHSMYGRNMHYPYHSHSASRHFDSPERWMDMSGYQMDMQGR<br>WMDNYGRYVNPFHHHMYGRNMFYPYGSHCNNRHMEHPERYMDMSGYQMDMQGRWMDTH<br>GRHCNPLGQMWHNRHGYYPGHPHGRNMFQPERWMDMSSYQMDMQGRWMDNYGRYVNPF<br>SHNYGRHMNYPGGHYNYHHGRYMNHPERQMDMSGYQMDMHGRWMDNQGRYIDNFDRNY<br>YDYHMY* |
| **A1 Y2EEEEE** | MNRYLNRQRLYNM**EEEEE**RNKYRGVMEPMSRMTMDFQGRYMDSQGRMVDPRYYDHYGR<br>MHDYDRYYGRSMFNQGHSMDSQRYGGWMDNPERYMDMSGYQMDMQGRWMDAQGRYNNP<br>FSQMWHSRQGHYPG**EEEEE**MSHHSMYGRNMHYPYHSHSASRHFDSPERWMDMSGYQMD<br>MQGRWMDNYGRYVNPFHHHMYGRNMFYPYGSHCNNRHMEHPERYMDMSGYQMDMQGRW<br>MDTHGRHCNPLGQMWHNRHGYYPGHPHGRNMFQPERWMDMSSYQMDMQGRWMDNYGRY |

| | |
|---|---|
| | VNPFSHNYGRHMNYPGGHYNYHHGRYMNHPERQMDMSGYQMDMHGRWMDNQGRYIDNF<br>DRNYYDYHMY* |
| **A2 WT**<br><br>UniProt Accession:<br>A0A088MI16 | MNRYMMRHRPM**Y**SNMYRTGRKYRGVMEPMSRMTMDFQGRYMDSQGRMVDPRYYEYGRC<br>HDYDRYNGRSMFNNGPYMDGQRYGGWMDFPERYMDMSGYQMDMHGRWMDSQGRYCNPM<br>GHSWSNRQGYYPGSNYGRNMFNPERYMDMSGYQMDMQGRWMDMGGRHVNPFSHSMYGR<br>NMFNPSYFSNRHMDNPERYMDMSGYQMDMQGRWMDTQGR**Y**MDP**S**MSNM**Y**DNYNYWY* |
| **A2 Y4E** | MNRYMMRHRPM**E**SNMYRTGRKYRGVMEPMSRMTMDFQGRYMDSQGRMVDPRYYEYGRC<br>HDYDRYNGRSMFNNGPYMDGQRYGGWMDFPERYMDMSGYQMDMHGRWMDSQGRYCNPM<br>GHSWSNRQGYYPGSNYGRNMFNPERYMDMSGYQMDMQGRWMDMGGRHVNPFSHSMYGR<br>NMFNPSYFSNRHMDNPERYMDMSGYQMDMQGRWMDTQGR**E**MDP**E**MSNM**E**DNYNYWY* |
| **A2 Y4EEE** | MNRYMMRHRPM**EEE**SNMYRTGRKYRGVMEPMSRMTMDFQGRYMDSQGRMVDPRYYEYG<br>RCHDYDRYNGRSMFNNGPYMDGQRYGGWMDFPERYMDMSGYQMDMHGRWMDSQGRYCN<br>PMGHSWSNRQGYYPGSNYGRNMFNPERYMDMSGYQMDMQGRWMDMGGRHVNPFSHSMY<br>GRNMFNPSYFSNRHMDNPERYMDMSGYQMDMQGRWMDTQGR**EEE**MDP**EEE**MSNM**EEE**D<br>NYNYWY* |
| **A2 Y4EEEEE** | MNRYMMRHRPM**EEEEE**SNMYRTGRKYRGVMEPMSRMTMDFQGRYMDSQGRMVDPRYYE<br>YGRCHDYDRYNGRSMFNNGPYMDGQRYGGWMDFPERYMDMSGYQMDMHGRWMDSQGRY<br>CNPMGHSWSNRQGYYPGSNYGRNMFNPERYMDMSGYQMDMQGRWMDMGGRHVNPFSHS<br>MYGRNMFNPSYFSNRHMDNPERYMDMSGYQMDMQGRWMDTQGR**EEEEE**MDP**EEEEE**MS<br>NM**EEEEE**DNYNYWY* |
| **B WT**<br><br>UniProt Accession:<br>A0A088MEC8 | MSSFMDPMHYDGMGMSHSKSGDFSHNCMRSFHKSQRDGMRRDIMGKSSKNRRFGNLME<br>PMSRMTMDFHGRLIDSQGRIVDPGHYFAMDDHYMENDRFLYPHDMLRNRHGMYGFMQG<br>DYGNNMHRGMFADGMYRDMHHSGMNPSSYMHGGSMQNRPMMYMQGRYLDDSYFMNYHD<br>PPVIVHSHYNDQEGRHQGMYDRHSDSYGSHRRHGDSHSMPRRPSESHSPQRRP**S**EGHI<br>IQVRPEGGSSRKTSRAQLFPDDKLTD**S**A* |
| **B S2E** | MSSFMDPMHYDGMGMSHSKSGDFSHNCMRSFHKSQRDGMRRDIMGKSSKNRRFGNLME<br>PMSRMTMDFHGRLIDSQGRIVDPGHYFAMDDHYMENDRFLYPHDMLRNRHGMYGFMQG<br>DYGNNMHRGMFADGMYRDMHHSGMNPSSYMHGGSMQNRPMMYMQGRYLDDSYFMNYHD<br>PPVIVHSHYNDQEGRHQGMYDRHSDSYGSHRRHGDSHSMPRRPSESHSPQRRP**E**EGHI<br>IQVRPEGGSSRKTSRAQLFPDDKLTD**E**A* |
| **B S2EEE** | MSSFMDPMHYDGMGMSHSKSGDFSHNCMRSFHKSQRDGMRRDIMGKSSKNRRFGNLME<br>PMSRMTMDFHGRLIDSQGRIVDPGHYFAMDDHYMENDRFLYPHDMLRNRHGMYGFMQG<br>DYGNNMHRGMFADGMYRDMHHSGMNPSSYMHGGSMQNRPMMYMQGRYLDDSYFMNYHD<br>PPVIVHSHYNDQEGRHQGMYDRHSDSYGSHRRHGDSHSMPRRPSESHSPQRRP**EEE**EG<br>HIIQVRPEGGSSRKTSRAQLFPDDKLTD**EEE**A* |
| **C**<br><br>UniProt Accession:<br>A0A088MRL7 | MNKHSSSHGMHGENYSRTGARGLHRGMEHESKSMYGKGRERSTDHGDMESSRHGMPGG<br>MNPGMYGGMPSGMPGGMYPGMYGGMPGFGVPQMMQCPPDMPRRYIDRHDRSMDMPYGR<br>YMDMPQGRYMSSQDRLMHMMHNRHLYGRMMDQGRMGEPMEGNMENRGRNMENYE* |

**Supplemental Table 2. Individual FRAP curve fitting data – homotypic condensates**

**Legend**

**# Cells**: Raw count of individual cell FRAPs measured. Count excludes measurements where fit $R^2$ < 0.70, extensive condensate movement laterally or out of the focal plane occurred during measurement, or other technical problems or errors.

**#Rec >15% / %Rec >15%**: Number and % of individual FRAP curves that have a normalized recovery at final data point ≥ 15%. Condensates that have <15% recovery are defined as dynamically arrested.

**$M_f$ Mean / $M_f$ S.D.**: Average mobile fraction and standard deviation from successful fits. For individual fits with normalized recovery < 15%, $M_f$ was estimated using the final data point (constructs with no cells >15% recovery, lines are in red.)

**$t_{1/2}$ Mean / $t_{1/2}$ S.D.:** Average recovery half-time and standard deviation from successful fit. For constructs for which no cells reached the selection criteria above, $t_{1/2}$ is listed as NA.

| Construct | Compartment | # Cells | # Rec >15% | % Rec > 15% | $M_f$ Mean | $M_f$ S.D. | $t_{1/2}$ Mean | $t_{1/2}$ S.D. |
|---|---|---|---|---|---|---|---|---|
| A1(WT)-eGFP | Nucleus | 31 | 19 | 61% | 0.27 | 0.24 | 80.49 | 71.51 |
| A1(WT)-eGFP | Cytoplasm | 25 | 0 | 0% | 0.01 | 0.01 | NA | NA |
| A1(LO)-eGFP | Nucleus | 27 | 25 | 93% | 0.31 | 0.13 | 53.04 | 38.49 |
| A1(LO-Extended)-eGFP | Nucleus | 31 | 29 | 94% | 0.31 | 0.20 | 52.87 | 48.39 |
| A1(RM)-eGFP | Nucleus | 28 | 28 | 100% | 0.92 | 0.07 | 4.78 | 1.92 |
| A1(RM)-mScarlet | Nucleus | 29 | 29 | 100% | 0.94 | 0.06 | 4.19 | 1.23 |
| A1(LnRM)-eGFP | Nucleus | 34 | 34 | 100% | 0.81 | 0.12 | 12.65 | 5.56 |
| A1(Y2EEE)-eGFP | Cytoplasm | 26 | 0 | 0% | 0.03 | 0.03 | NA | NA |
| A1(Y2EEEEE)-eGFP | Nucleus | 29 | 22 | 76% | 0.31 | 0.23 | 65.51 | 59.64 |
| A1(Y2EEEEE)-eGFP | Cytoplasm | 28 | 0 | 0% | 0.03 | 0.02 | NA | NA |
| A2(WT)-eGFP | Nucleus | 26 | 22 | 85% | 0.61 | 0.30 | 106.2 | 51.38 |
| A2(WT)-mScarlet | Nucleus | 30 | 10 | 33% | 0.22 | 0.23 | 125.1 | 109.6 |
| A2(WT)-eGFP | Cytoplasm | 30 | 0 | 0% | 0.04 | 0.02 | NA | NA |
| A2(Y4E)-eGFP | Nucleus | 33 | 27 | 82% | 0.56 | 0.30 | 78.94 | 50.46 |
| A2(Y4EEE)-eGFP | Nucleus | 31 | 31 | 100% | 0.82 | 0.17 | 46.01 | 32.25 |
| A2(Y4EEEEE)-eGFP | Nucleus | 41 | 41 | 100% | 0.80 | 0.10 | 14.88 | 5.74 |
| B(WT)-eGFP | Nucleus | 24 | 24 | 100% | 0.85 | 0.08 | 6.74 | 2.87 |
| B(WT)-mScarlet | Nucleus | 35 | 35 | 100% | 0.88 | 0.08 | 7.13 | 2.01 |
| B(WT)-mScarlet | Cytoplasm | 29 | 29 | 100% | 0.74 | 0.16 | 7.31 | 5.13 |
| B(S2E)-eGFP | Nucleus | 25 | 25 | 100% | 0.90 | 0.10 | 7.08 | 4.10 |
| B(S2EEE)-eGFP | Nucleus | 30 | 30 | 100% | 0.90 | 0.07 | 5.29 | 1.89 |
| B(S2EEE)-mScarlet | Nucleus | 28 | 28 | 100% | 0.93 | 0.06 | 7.97 | 3.19 |
| C(WT)-eGFP | Nucleus | 35 | 35 | 100% | 0.85 | 0.11 | 12.29 | 5.63 |
| C(WT)-mScarlet | Nucleus | 27 | 26 | 96% | 0.93 | 0.07 | 3.60 | 1.18 |

**Supplemental Table 3. Individual FRAP curve fitting data – heterotypic condensates**

**Legend**

**# Cells**: Raw count of individual cell FRAPs measured. Count excludes measurements where fit $R^2$ < 0.70, extensive condensate movement laterally or out of the focal plane occurred during measurement, or other technical problems or errors.

**#Rec >15% / %Rec >15%**: Number and % of individual FRAP curves that have a normalized recovery at final data point ≥ 15%. Condensates that have <15% recovery are defined as dynamically arrested.

**$M_f$ Mean / $M_f$ S.D.**: Average mobile fraction and standard deviation from successful fits. For individual fits with normalized recovery < 15%, $M_f$ was estimated using the final data point (constructs with no cells >15% recovery, lines are in red.)

**$t_{1/2}$ Mean / $t_{1/2}$ S.D.:** Average recovery half-time and standard deviation from successful fit. For constructs for which no cells reached the selection criteria above, $t_{1/2}$ is listed as NA.

| Construct | Compartment | # Cells | # Rec >15% | % Rec >15% | $M_f$ Mean | $M_f$ S.D. | $t_{1/2}$ Mean | $t_{1/2}$ S.D. |
|---|---|---|---|---|---|---|---|---|
| **A1(LO)-eGFP*** : A1(RM)-mScarlet | Nucleus | 34 | 34 | 100% | 0.60 | 0.23 | 53.48 | 27.09 |
| A1(LO)-eGFP **: A1(RM)-mScarlet*** | Nucleus | 34 | 33 | 97% | 0.83 | 0.11 | 5.74 | 2.26 |
| **A1(WT)-eGFP*** : A2(WT)-mScarlet | Nucleus | 31 | 11 | 35% | 0.23 | 0.27 | 140.6 | 155.0 |
| A1(WT)-eGFP : **A2(WT)-mScarlet*** | Nucleus | 31 | 5 | 16% | 0.19 | 0.27 | 221.9 | 143.3 |
| **A1(WT)-eGFP*** : B(WT)-mScarlet | Nucleus | 31 | 27 | 87% | 0.71 | 0.30 | 58.76 | 51.05 |
| A1(WT)-eGFP : **B(WT)-mScarlet*** | Nucleus | 32 | 32 | 100% | 0.87 | 0.11 | 11.55 | 3.70 |
| **A1(WT)-eGFP*** : B(WT)-mScarlet | Cytoplasm | 24 | 0 | 0% | 0.05 | 0.03 | NA | NA |
| A1(WT)-eGFP : **B(WT)-mScarlet*** | Cytoplasm | 23 | 23 | 100% | 0.72 | 0.13 | 10.66 | 4.97 |
| **A1(Y2EEEEE)-eGFP*** : B(WT)-mScarlet | Nucleus | 32 | 22 | 69% | 0.47 | 0.35 | 77.59 | 88.40 |
| A1(Y2EEEEE)-eGFP : **B(WT)-mScarlet*** | Nucleus | 32 | 32 | 100% | 0.84 | 0.09 | 12.78 | 6.75 |
| **A1(Y2EEEEE)-eGFP*** : B(WT)-mScarlet | Cytoplasm | 25 | 3 | 12% | 0.06 | 0.06 | 36.38 | 24.24 |
| A1(Y2EEEEE)-eGFP : **B(WT)-mScarlet*** | Cytoplasm | 25 | 25 | 100% | 0.72 | 0.15 | 23.16 | 18.25 |
| **A2(WT)-eGFP*** : B(WT)-mScarlet | Nucleus | 32 | 30 | 94% | 0.68 | 0.27 | 51.23 | 42.22 |
| A2(WT)-eGFP : **B(WT)-mScarlet*** | Nucleus | 32 | 32 | 100% | 0.82 | 0.09 | 10.95 | 3.90 |
| **A2(WT)-eGFP*** : C(WT)-mScarlet | Nucleus | 29 | 29 | 100% | 0.76 | 0.21 | 71.53 | 38.80 |
| A2(WT)-eGFP : **C(WT)-mScarlet*** | Nucleus | 29 | 28 | 97% | 0.80 | 0.11 | 5.81 | 4.39 |
| **A2(WT)-eGFP*** : B(S2EEE)-mScarlet | Nucleus | 33 | 32 | 97% | 0.72 | 0.24 | 72.79 | 59.25 |
| A2(WT)-eGFP : **B(S2EEE)-mScarlet*** | Nucleus | 33 | 33 | 100% | 0.78 | 0.10 | 9.67 | 4.36 |
| **A2(Y4EEE)-eGFP*** : B(WT)-mScarlet | Nucleus | 30 | 30 | 100% | 0.86 | 0.14 | 25.39 | 14.37 |
| A2(Y4EEE)-eGFP : **B(WT)-mScarlet*** | Nucleus | 29 | 29 | 100% | 0.86 | 0.12 | 11.80 | 5.26 |

**Supplemental Figure 1. Complete amino acid compositional analysis of linker and domain regions of *D. opalescens* reflectins.** Relatively enriched amino acids are highlighted in red, relatively depleted amino acids are highlighted in blue.

% Amino Acid

| | A1-WT | A1-RM | A1-LnRM | A1-LO | A2-WT | A2-RM | A2-LO | B-WT | B-RM | B-LO | C-WT | C-RM | C-LO | C-GMXX | C-LO-GMXX |
|---|---|---|---|---|---|---|---|---|---|---|---|---|---|---|---|
| #AA | 350 | 166 | 187 | 184 | 230 | 109 | 121 | 260 | 25 | 235 | 170 | 28 | 110 | 32 | 142 |
| A | 0.6 | 0.6 | 0.5 | 0.5 | 0.0 | 0.0 | 0.0 | 1.5 | 0.0 | 1.7 | 0.6 | 0.0 | 0.9 | 0.0 | 0.7 |
| C | 0.6 | 0.6 | 0.5 | 0.5 | 0.9 | 0.9 | 0.8 | 0.4 | 0.0 | 0.4 | 0.6 | 0.0 | 0.9 | 0.0 | 0.7 |
| D | 7.7 | 11.4 | 10.2 | 4.3 | 8.3 | 11.9 | 5.0 | 9.6 | 12.0 | 9.4 | 5.3 | 14.3 | 4.5 | 0.0 | 3.5 |
| E | 2.0 | 3.6 | 3.2 | 0.5 | 2.2 | 3.7 | 0.8 | 2.3 | 4.0 | 2.1 | 5.9 | 0.0 | 9.1 | 0.0 | 7.0 |
| F | 2.6 | 1.2 | 1.1 | 3.8 | 3.0 | 0.9 | 5.0 | 4.2 | 4.0 | 4.3 | 0.6 | 0.0 | 0.9 | 0.0 | 0.7 |
| G | 10.0 | 9.6 | 9.1 | 10.3 | 10.9 | 11.0 | 10.7 | 10.0 | 8.0 | 10.2 | 17.1 | 7.1 | 12.7 | 40.6 | 19.0 |
| H | 8.9 | 1.8 | 1.6 | 15.2 | 3.0 | 1.8 | 4.1 | 8.8 | 4.0 | 9.4 | 6.5 | 3.6 | 9.1 | 0.0 | 7.0 |
| I | 0.3 | 0.6 | 0.5 | 0.0 | 0.0 | 0.0 | 0.0 | 2.3 | 8.0 | 1.7 | 0.6 | 3.6 | 0.0 | 0.0 | 0.0 |
| K | 0.3 | 0.0 | 0.5 | 0.5 | 0.4 | 0.0 | 0.8 | 2.3 | 0.0 | 2.6 | 1.8 | 0.0 | 2.7 | 0.0 | 2.1 |
| L | 0.9 | 0.0 | 1.1 | 1.6 | 0.0 | 0.0 | 0.0 | 2.7 | 4.0 | 2.6 | 1.8 | 0.0 | 2.7 | 0.0 | 2.1 |
| M | 14.0 | 18.7 | 17.6 | 9.8 | 16.5 | 21.1 | 12.4 | 11.5 | 16.0 | 11.1 | 17.6 | 17.9 | 15.5 | 25.0 | 17.6 |
| N | 7.4 | 5.4 | 7.0 | 9.2 | 7.8 | 1.8 | 13.2 | 4.2 | 0.0 | 4.7 | 4.7 | 0.0 | 6.4 | 3.1 | 5.6 |
| P | 4.9 | 6.6 | 5.9 | 3.3 | 5.2 | 7.3 | 3.3 | 5.4 | 8.0 | 5.1 | 7.6 | 10.7 | 3.6 | 18.8 | 7.0 |
| Q | 6.0 | 8.4 | 8.0 | 3.8 | 4.8 | 8.3 | 1.7 | 3.8 | 4.0 | 3.8 | 2.9 | 3.6 | 3.6 | 0.0 | 2.8 |
| R | 10.9 | 10.8 | 12.3 | 10.9 | 11.7 | 11.0 | 12.4 | 10.4 | 12.0 | 10.2 | 10.6 | 21.4 | 10.9 | 0.0 | 8.5 |
| S | 6.0 | 4.8 | 4.3 | 7.1 | 7.4 | 5.5 | 9.1 | 11.9 | 8.0 | 12.3 | 7.6 | 3.6 | 10.0 | 3.1 | 8.5 |
| T | 0.6 | 1.2 | 1.1 | 0.0 | 1.3 | 1.8 | 0.8 | 1.2 | 4.0 | 0.9 | 1.2 | 0.0 | 1.8 | 0.0 | 1.4 |
| V | 1.1 | 1.8 | 2.1 | 0.5 | 1.3 | 1.8 | 0.8 | 1.5 | 4.0 | 1.3 | 0.6 | 0.0 | 0.9 | 0.0 | 0.7 |
| W | 2.9 | 4.2 | 3.7 | 1.6 | 2.6 | 2.8 | 2.5 | 0.0 | 0.0 | 0.0 | 0.0 | 0.0 | 0.0 | 0.0 | 0.0 |
| Y | 12.6 | 8.4 | 9.6 | 16.3 | 12.6 | 8.3 | 16.5 | 5.8 | 0.0 | 6.4 | 6.5 | 14.3 | 3.6 | 9.4 | 4.9 |

**Supplemental Figure 2. mScarlet-tagged reflectins demonstrate similar condensates to eGFP-tagged ones by confocal microscopy.** Confocal images of mScarlet-labeled reflectin WT isoforms. Scale bar = 10 μm.

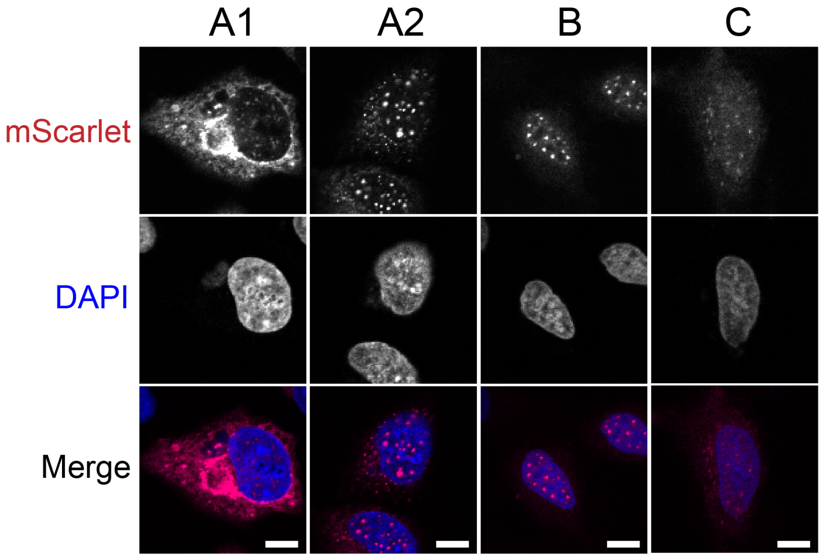

**Supplemental Figure 3. Western blotting confirms successful expression of reflectin constructs in HeLa cells.** Western blots of eGFP-tagged A) WT and B) mutant reflectins used in this study. C) mScarlet-tagged reflectins used in this study.

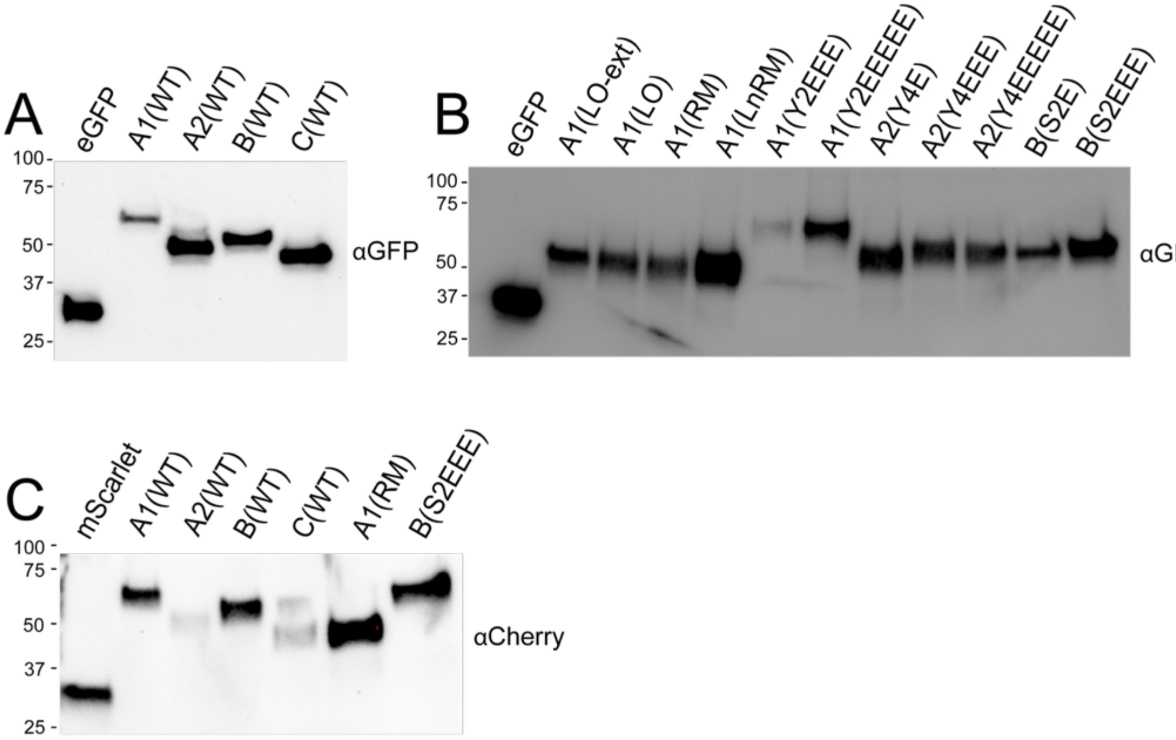

**Supplemental Figure 4. Reflectin proteins are not strongly phosphorylated in HeLa cells.** A) ProQ-stained blot of HeLa cells expressing different WT reflectins, and B) western blot of the same lanes. The positive control was a protein standard containing phosphorylated Ovalbumin protein at 45kDa (Cat No. P6649, Thermofisher)

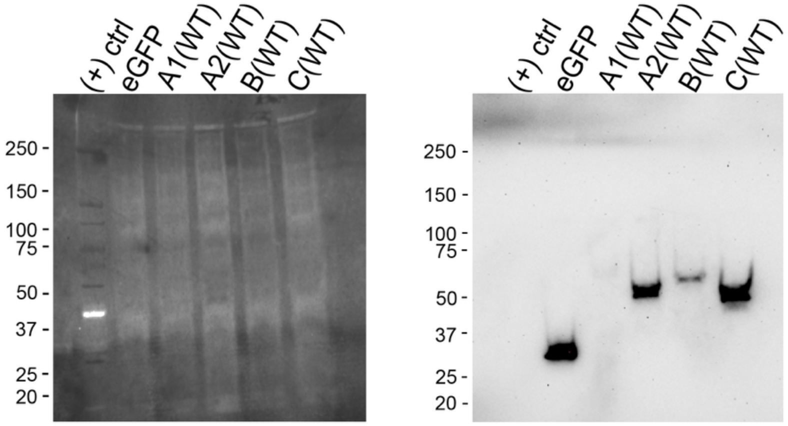

**Supplemental Figure 5. Reflectin B is insensitive to fluorescent protein tag and displays similar dynamics in both the nuclear and cytoplasmic compartments.** A) Representative FRAP curves of reflectin B across the nuclear and cytoplasmic compartments. Both eGFP- and mScarlet tagged B show similar dynamics. B) $M_f$ and C) $t_{1/2}$ distributions after fitting of each individual cell fluorescence recovery curve ($n$ = 3 biological replicates). B-eGFP and B-mScarlet data reproduced from Fig. 3 for comparison.

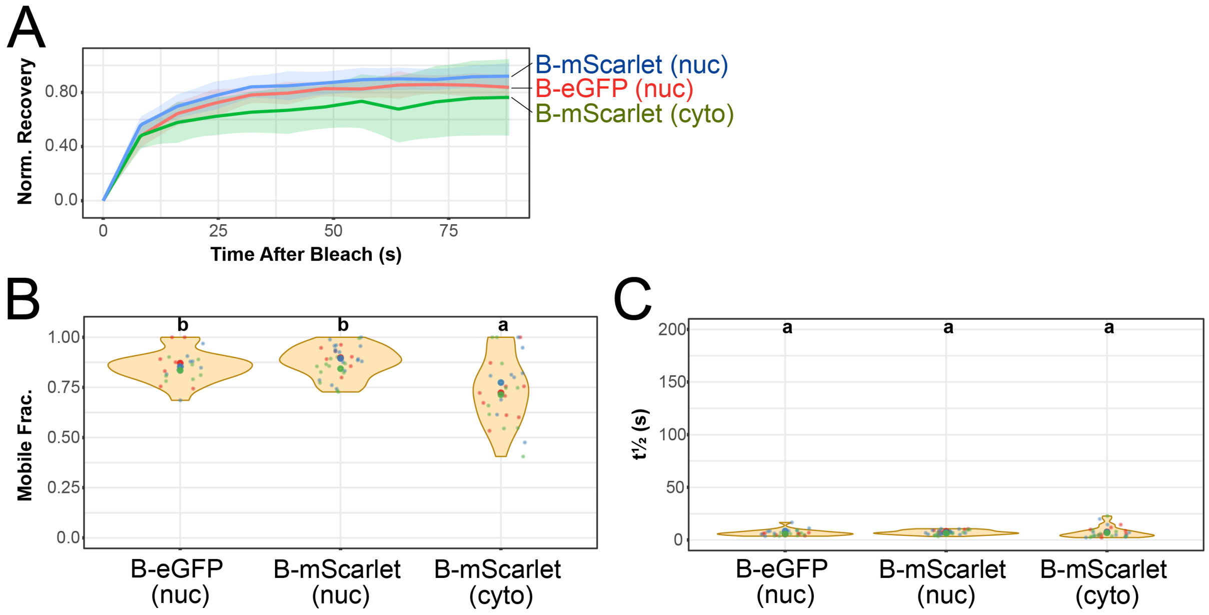

**Supplemental Figure 6. Two A1 linker-only variants both display similar, low fluorescence recovery.** A) Representative confocal images of eGFP-tagged A1 LO-Extended (A1 LO-Ext), compared to A1 LO. Scale bar = 10 µm. B) Representative FRAP curves comparing A1 LO and A1 LO-Extended. B) $M_f$ and C) $t_{1/2}$ distributions after fitting of each individual cell fluorescence recovery curve ($n$ = 3 biological replicates). A1 LO data reproduced from Fig. 4 for comparison.

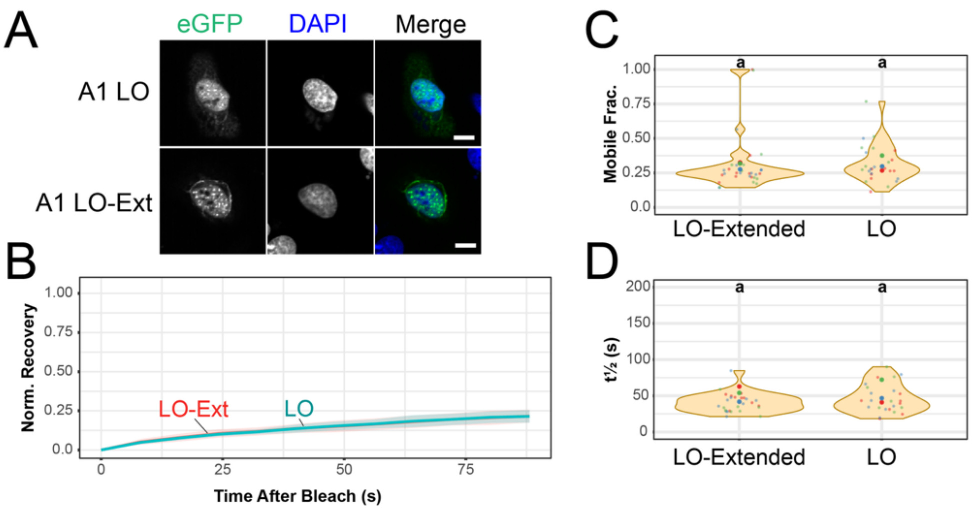

**Supplemental Figure 7. Reflectin A1, A2, and B phosphomimetic mutants form condensates.** Confocal images of A) reflectin A1, B) reflectin A2, and C) reflectin B phosphomimetic mutants. Scale bar = 10 μm.

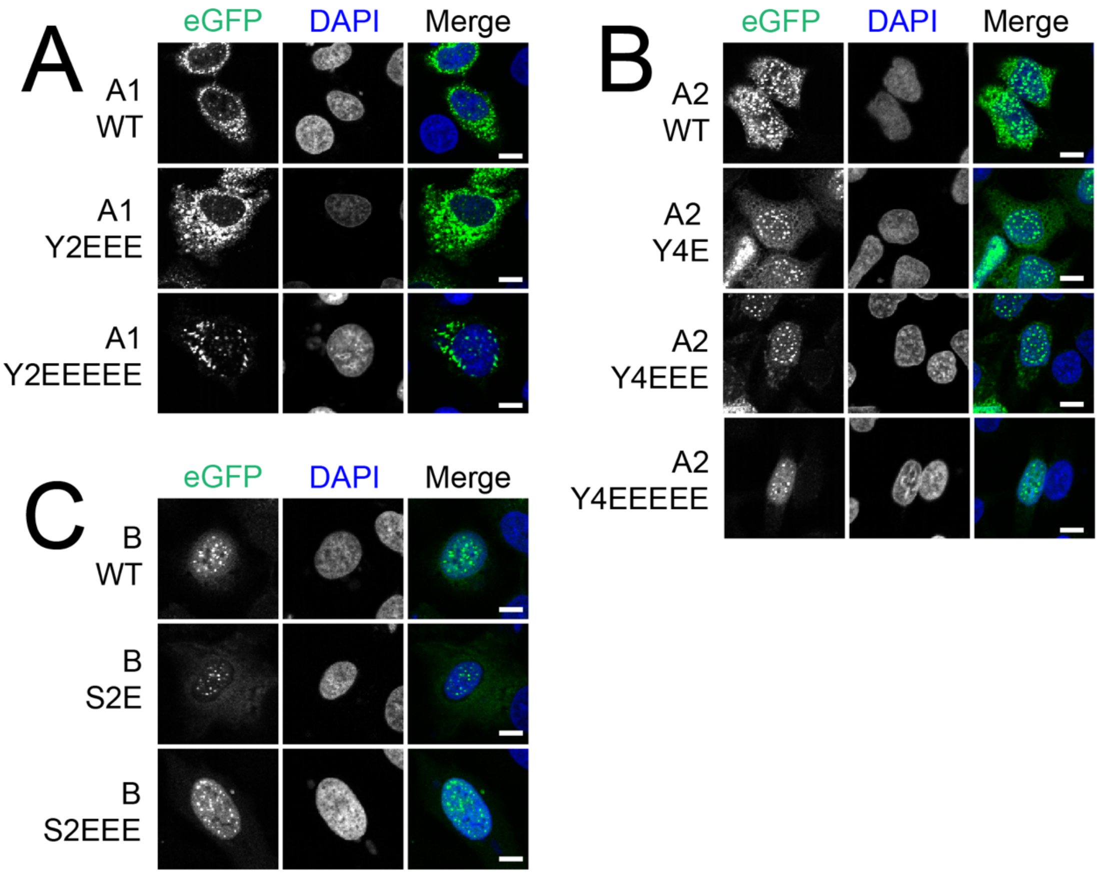

**Supplemental Figure 8. Heterotypic condensates of A1 and A2 do not display increased fluidity.** **A)** Representative FRAP curves of reflectin A1 (left) or A2 (right) in heterotypic A1:A2 condensates. B) $M_f$ and C) $t_{1/2}$ distributions after fitting of all individual cells with final normalized recovery > 15% (11/31 cells for A1, 5/31 cells for A2, from *n* = 3 biological replicates). Means of individual biological replicates are shown as larger, centered dots. CLD letters at the top distinguish statistically significant differences between groups within each panel. A1-eGFP and A2-eGFP data reproduced from Fig. 3 for comparison.

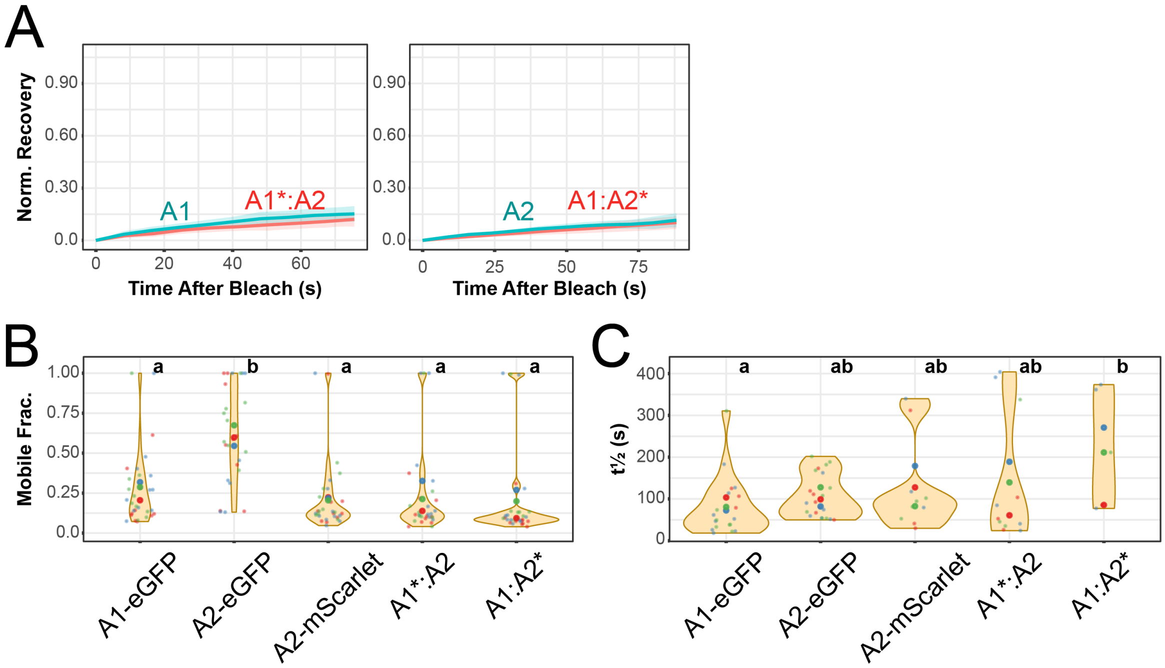

**Supplemental Figure 9. Heterotypic condensates of A2 with B or C do not substantially alter the fluidity of the B or C component.** A) Representative FRAP curves of reflectin B in heterotypic condensates across nuclear and cytoplasmic compartments. B, C) $M_f$ and $t_{1/2}$ distributions after fitting of the B or D,E) C component from all individual cells from $n$ = 3 biological replicates. Means of individual biological replicates are shown as larger, centered dots. CLD letters at the top distinguish statistically significant differences between groups within each panel. B-mScarlet and A1:B* data reproduced from Fig 3 and 6, respectively, for comparison.

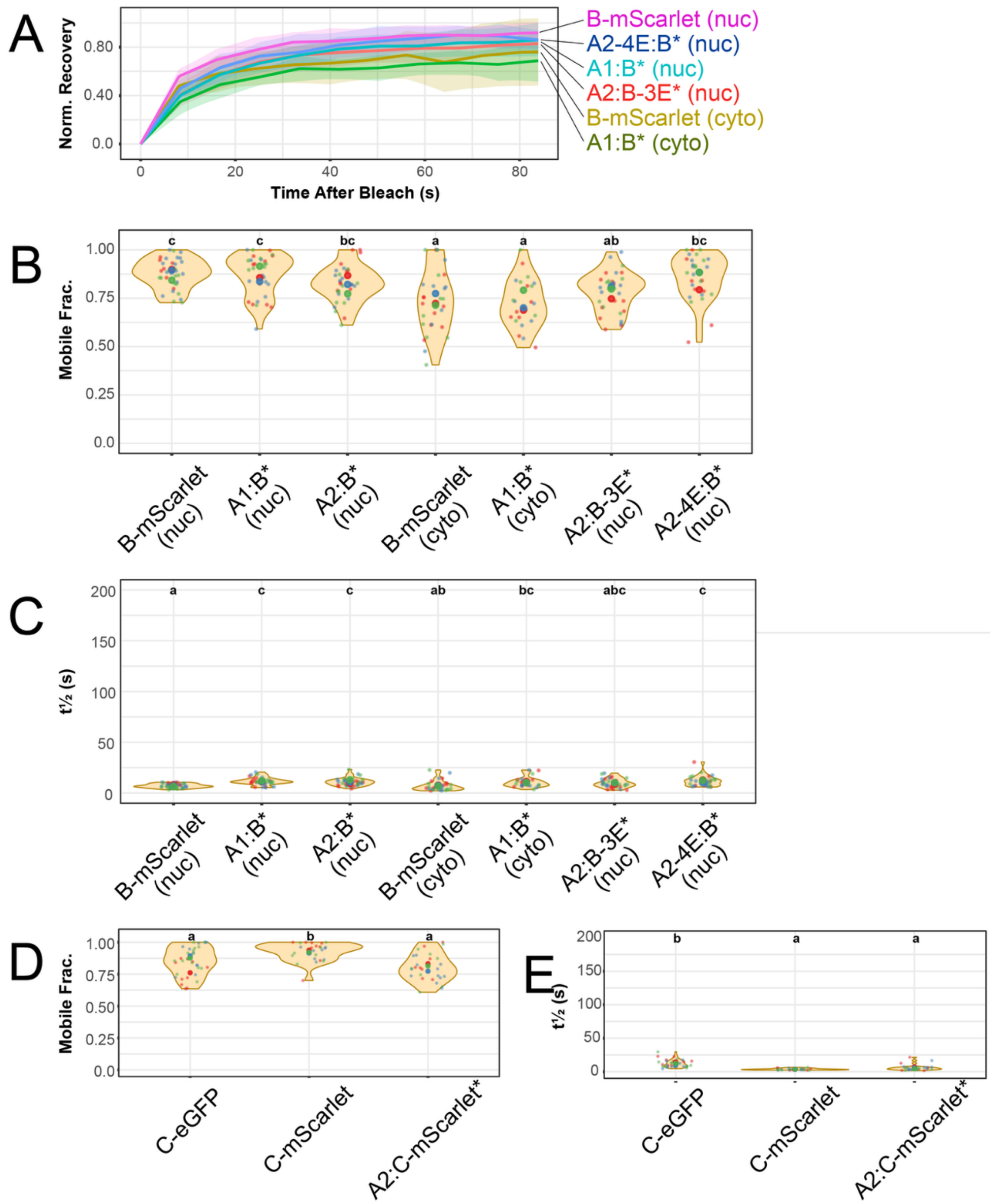
